# Prediction and information integration determine subtle anticipatory fixation biases

**DOI:** 10.1101/252809

**Authors:** Giuseppe Notaro, Wieske van Zoest, David Melcher, Uri Hasson

**Affiliations:** Center for Mind/Brain Sciences (CIMeC), The University of Trento, Italy; Center for Practical Wisdom, The University of Chicago, USA

## Abstract

A core question underlying neurobiological and computational models of behavior is how individuals learn environmental statistics and use them for making predictions. Treatment of this issue largely relies on reactive paradigms, where inferences about predictive processes are derived by modeling responses to stimuli that vary in likelihood. Here we deployed a novel proactive oculomotor metric to determine how input statistics impact anticipatory behavior, decoupled from stimulus-response. We implemented transition constraints between target locations, and quantified a subtle fixation bias (FB) discernible while individuals fixated a screen center awaiting target presentation. We show that FB is informative with respect the input statistics, reflects learning at different temporal scales, predicts saccade latencies on a trial level, and can be linked to fundamental oculomotor metrics. We also present an extension of this approach to a more complex paradigm. Our work demonstrates how learning impacts strictly predictive processes and presents a novel direction for studying learning and prediction.

## Introduction

Construction of precise predictions about future events optimizes perception (e.g., Auksztulewicz et al., 2017; Kok et al., 2012; Rohenkol et al., 2014) and selection of goal-directed action (Friston et al., 2006). Understanding how individuals acquire statistical knowledge about their environment, and whether they capitalize on it for making predictions are two questions at the core of current computational and neurobiological investigations (e.g., Clark, 2013; Emberson et al., 2015; Glascher & Buchel, 2005; Harrison et al., 2011; Krogh et al., 2012; Schapiro & Turk-Browne, 2015; Vossel et al., 2014). Statistical learning can be inferred from the analysis of responses to stimuli varying in predictability, since individuals respond more quickly and accurately when stimuli match their expectation (e.g., den Ouden et al., 2010). This idea also informs current experimental approaches where manual (Siegelman et al., 2017) or oculomotor responses (e.g., Aslin, 2014; Kidd et al., 2012; Vossel et al., 2014; Marcus et al., 2006) are treated as indices of statistical learning.

Yet, drawing conclusions about the very existence of predictive processes and their properties from stimulus-linked responses is accompanied by several complications. Responses are co-determined by multiple factors which certainly include prior knowledge and prediction, but also attentional orientation, low-level perception, evidence-accumulation, surprise, belief-updating, and decisions (see Bar et al., 2006; Grossberg, 1987; O’Reilly et al., 2013; Vossel et al., 2014). Each of these processes, and their interactions, can be impacted in different ways by prior predictions and noise. These complexities constitute an interesting challenge for learning and decision models that are precisely interested in how prior knowledge interacts with response-related elements such as stimulus surprise and updating of priors (e.g., Glascher et al, 2010), or how control mechanisms contribute to behavior (e.g., Jiang et al., 2014). However, this multi-factorial nature of responses complicates their use for understanding whether and how predictions arise and what sorts of learning processes specifically govern anticipatory processes in and of themselves. Advances in data modeling have certainly been made towards disambiguating between stimulus expectation (‘prior state‘, or ‘start point‘), processing of stimulus features (e.g., Carpenter & William, 1995; Reddi & Carpenter 2000), and updating of prior knowledge (e.g., Brodersen et al., 2008; Glaze et al., 2015; O’Reilly et al., 2013; Vossel et al., 2014). Nonetheless, even precise estimation of model parameters linked to prior-knowledge cannot directly speak to how this knowledge is linked to online processing: whether by supporting proactive predictions prior to stimulus presentation, or alternatively, via reactive, backward-looking integrative processes that are initiated after stimulus presentation (e.g., Grossberg, 1987). To determine whether predictions take place and model the learning trajectories that govern them, it is therefore essential to isolate and study prediction-related behavioral signatures that are identifiable prior to stimulus presentation.

The temporal constants that mediate predictive processes are of particular interest. It is known that stimulus responses are strongly impacted by stimulus history in the very recent past (e.g., Barton et al., 2006; Marcos et al., 2013), which is compatible with both error-driven learning (e.g., Rescorla & Wagner, 1972) or sequential effect models (Yu & Cohen, 2008).

An exponential decay of the impact of the previous stimuli has been confirmed by neuroimaging and behavioral studies (e.g., Bornstein & Daw, 2012; Harrison et al., 2011). Kim et al. (2017) manipulated the first-order transition dependency between target locations and reported that saccade latencies were related to the prior probability of making the specific saccade (see also Farrell et al., 2010); when modeled via LATER (Carpenter & Williams, 1995), this was reflected in sequential-dependent updates of the starting point of a rise-to-threshold saccade generation process. Given these well replicated findings it is essential to determine whether predictive behaviors show different sensitivity to events in the recent past than the responses themselves.

To understand whether and how statistical learning impacts predictive behavior, we developed an overt behavioral measure that reflects learning and prediction, but that is dissociated from responses to stimuli. In brief, we presented participants with series of visual trials that licensed predictions about the location of the next item, and we quantified gaze location during those parts of the trial *prior* to target presentation, while participants fixated the screen center. We used this fixation-bias measure to address several questions pertaining to predictive processes. First, do individuals at all routinely engage in proactive predictive behavior? Second, is predictive behavior determined solely by the distribution of events in the very recent past (as suggested e.g., by Harrison et al., 2011) or are predictive behaviors determined by learning that jointly occurs over both short and long temporal scales (as suggested, e.g., by Bornstein & Daw, 2012)? Third we determined whether fixation biases provide more information about the statistical structure of the environment than do stimulus-linked responses (in this case, saccade latencies to targets), and relatedly, whether fixation bias is more sensitive to trial history. Fourth, and finally, to understand the relation between anticipatory processes and responses, we determined the trial-by-trial correlation between fixation biases and saccade latencies.

We report two studies. Experiment 1 is foundational and presents the general paradigm for the case of a two-state Markov process, as well as associated methods and results. Experiment 2 is an extension that shows how anticipatory fixation biases can be studied in more complex contexts by using steady-state analysis methods that utilize frequency-space analysis to describe inter-trial patterns in anticipatory gaze location.

Anticipating the detailed results reported below, the statistical structure of the visual series induced subtle but highly robust fixation biases prior to target presentation, below 1° eccentricity from fixation center. We find that *i*) fundamental properties of statistical learning on multiple temporal scales can be read off from this purely anticipatory gaze behavior while participants are at fixation, and *ii*) that the information this anticipatory behavior provides about the inputs’ statistical structure and learning dynamics differs from and exceeds that provided by saccade latencies.

## Methods

### Participants

Twenty-one volunteers participated in the study. (Mean *Age* = 23.8 ± 0.9; SEM is measure of spread throughout unless noted otherwise). They were recruited from the local student population, and reimbursed 20 Euro for their time. The institutional Ethical Review Board approved the study. The sample size was predetermined based on a pilot study with a similar design but that used images of real-life objects rather than abstract shapes (*N* = 20).

### Design

Participants observed multiple series of 100 trials each. Each trial consisted of a fixation symbol appearing at center, followed by a target that appeared to the right or left of center (Figure 1). The design consisted of one factor with two levels – high or low probability of return to the screen side of the last target. Specifically, in one condition, the probability of returning to the same side (probability of return) was 70% and in the other condition it was 30% (*pret*70, *pret*30 respectively). There were 10 series in each condition. The transition probabilities were fixed (stationary) within each series. While the transition probabilities were experimentally manipulated, the proportion of presentations on the left and right screen sides were identical and set at 50% in both conditions. Thus, any differences in behavior could only be attributed to differences in transition structure. To compare learning indices for the first and second half of each series, we constructed the series so that the intended transition constraint and screen-side frequencies were exactly maintained across trials 1–50 and trials 51–100.

**Figure 1.**
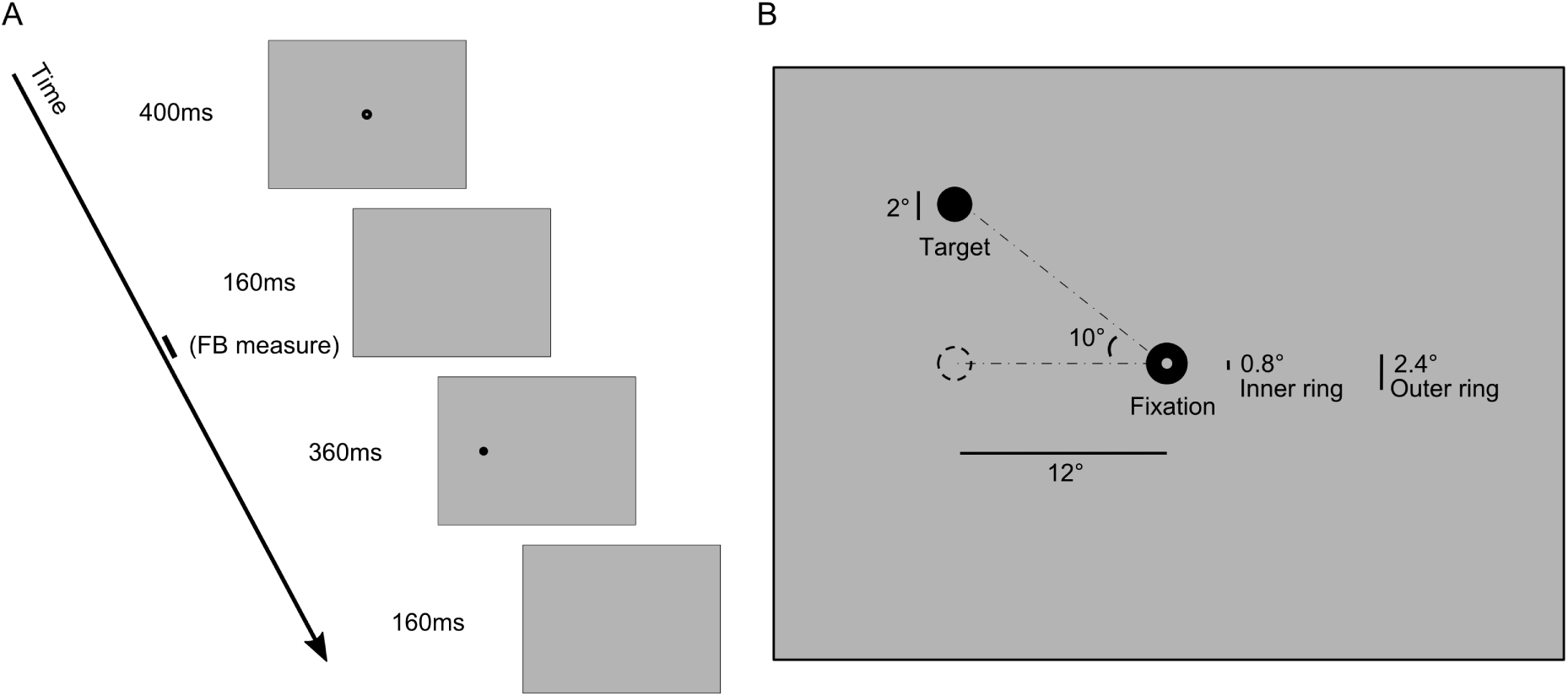
Trial structure and fixation locations in Experiment 1 (presented to scale). Panel A. Trial Timing. Fixation Bias was defined at the mean gaze location during the last 10 millisecond of the blank screen that followed the fixation symbol and that preceded the target. **Panel B**: Spatial features of fixation and targets. Targets were positioned on an invisible arch that extended 10° above and below the fixation symbol, at 12° eccentricity. Specific location on the arch was always determined randomly. The fixation symbol consisted of an inner ring (*radius* = 0.4°) within an outer ring (*radius* = 1.2°).

To each 100-trial series we appended 20 trials whose screen side was randomly determined. These were included to evaluate the impact of the prior series’ transition structure on responses to random trials (a transfer measure) and to aid clearing memory of the current stochastic process before beginning the next series.

## Procedure

### Eye-tracking

Stimuli were displayed on a CRT display (Diamond pro 2070SB, Mitsubishi Electric Corporation, Tokyo, Japan) with a spatial resolution of 1280 × 1024 pixels, set to a refresh rate of 75Hz. All the experimental software was generated using Matlab^TM^ and the Psychophysics Toolbox extensions (Brainard, 1997). Participants’ eyes were set at the same height as the screen center and at a distance of 58 cm. Eye position signals were recorded by a separate computer with a head-mounted, video-based eye tracker (*Eyelink 1000* Tower mount, SR Research Ltd, Mississauga, Canada) and were sampled monocularly at 1000 Hz. We performed a nine-point calibration procedure during which the eye-tracker calculated a mapping between sensor and display positions. To increase the accuracy of this mapping we performed calibration only in a display region that was slightly larger than the area used in the study (960 × 718 pixels around the center). We performed calibration after each break. Before beginning the experiment we identified each participant‘s dominant eye using the Dolman method.

### Trial structure

Participants were instructed to saccade to a target presented after the fixation symbol disappeared. The timeline of each trial (see Figure 1) was as follows. a fixation symbol appeared for 400ms; a post-fixation blank screen for 160ms; the target for 360ms; and a post-target blank screen for 160ms. The fixation symbol consisted of an inner gray circle with radius of 0.4° (same color as background) within an outer black circle with radius of 1.2°. We chose this fixation symbol as it has been shown to allow some variance in eye movement during fixation (Thaler, Schutz, Goodale, & Gegenfurtner, 2013). Targets were black circles with 1° radius that appeared to the left or right of the screen center, at 12° eccentricity (Figure 1B). The target-centers were located on a virtual (invisible) arch extending 10° vertically above and below the horizontal midline. On any given trial, the target’s specific position on the arc was set randomly. Participants could therefore anticipate the screen side of the next target but not its exact location. The specific instructions given were to saccade rapidly to the target and fixation symbol when they appeared.

### Instructions and training

To maintain participants’ alertness, we included catch trials in the form of target symbols with a white line through them. These appeared every 16–20 trials following a uniform distribution. Participants were told that catch trials would appear infrequently and that they were to press the mouse button when they saw those. Following each series, participants were presented with performance indicators for the last series, which included the number of hit targets, hit fixation symbols, correct catch trials and number of eye blinks, as well as their overall mean performance to that point. This was done to motivate participants to perform better and further buffer between subsequent series.

Before beginning the study proper, participants underwent training where they viewed series of 20 trials each, until they were comfortable with the procedure (typically within 2–7 sessions). In the training series there were no transition constraints (probability of return = 50%). During training we provided audio feedback in real-time: we provided positive audio feedback whenever participants’ gaze hit the target or the fixation circle within 200ms from appearance and with a maximum deviation of 1 ° from their border, and whenever participants correctly responded to catch trials with a mouse click. We provided negative audio feedback whenever participants failed to hit the target, failed to respond to catch trials, or blinked. A summary of the positive and negative scores was presented at the end of each training session.

## Analysis

### Saccade classification

During the study, participants performed large saccades to reach the targets from fixation circles. They also made many smaller saccades to adjust their gaze, typically during fixation. To detect saccades varying across this range, we applied a method (Nyström & Holmqvist, 2010) that detects saccades by determining speed thresholds adaptively. We defined saccade onset time as the time of the first minimum preceding a saccade peak. We defined saccade offset time as the last minimum after a peak, exceeding a threshold determined adaptively. When a saccade was followed by an assessment oscillation (glissade), the time of saccade offset was considered as the end of the glissade.

### Trial selection

We defined valid trials as ones where participants made saccades to both the fixation symbol and subsequent target, within a tolerance of 3° from their edge on the horizontal axis. To exclude anticipatory saccade we considered only trials where saccade latencies exceeded 80ms (Fischer & Ramsperger, 1984). Catch trials and the trial immediately following them were excluded from the analysis: these valid trials accounted for 87 ± 2% of the data, indicating good compliance with instructions.

We defined *Gaze Bias* as the mean gaze location measured during the last 10ms of the post-fixation blank, prior to target presentation. We then defined *Fixation Bias (FB)* as Gaze Bias in X direction, coded as positive if to the side of the last target and negative otherwise. On any given trial we defined an FB measurement as valid if three conditions held: *i*) there were no saccades or micro-saccades within this 10ms period, *ii*) both the target of the prior trial and current trial were correctly saccaded to, and *iii*) gaze was within a threshold radius *rad_FB_* = 3° from screen center. This last constraint was included solely to reduce the impact of eye position on saccade latency, as excessive positioning away from center could translate into faster arrival at target. We verified (see *Additional Information*) that the choice of *rad_FB_* did not alter the main findings for the Fixation Bias analysis. Following this FB definition, we further restricted our analyses to valid trials that were preceded by a correct fixation to the prior target. These trials accounted for 65 ± 2% of all trials.

### Impact of recent trials

To determine the impact of previous trials on current oculomotor behavior (as captured by FB) we defined two kinds of trials; *returns* which were trials where the screen-side of the last-presented target was the same as the one that preceded it, and *alternation* where the screen-side of the last-presented target was the opposite of the one preceding it. In this schema, FB quantifies the impact of the last transition (categorized as return or alternation) on anticipatory oculomotor behavior. Saccade latencies were analyzed according to the same schema.

In a separate analysis we modeled the impact of each of the last 6 transitions on current oculomotor behavior. We used a regression model in which dummy variables coded the status of each of the last six transitions as a return or alternation. This approach has been successfully used in prior work on statistical learning of transition probabilities (e.g., Bornstein & Daw, 2012). The complete regression model is presented in Equation 1, where S = 1 if the trial is a return, and 0 if alternation. This information is coded for each of the last *k* transitions (*k* = 6).

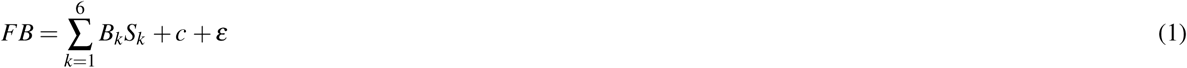

In this model, positive coefficients for any of the regressors *β*_1_ to *β*_6_ indicate that a return at lag *k* was associated with increased FB. Negative coefficients indicate reduced FB. The intercept *c* is the expected FB for 6 consecutive alternations, and is not further considered. When analyzing FB data, we fit these regression models to each participant, predicting the current FB value separately for the pret70 and pret30 conditions.

For saccade latencies (*SL*), we similarly fit regressions separately for the two conditions, but constructed separate models for return and alternation saccades. This is because return saccades are strongly impacted by inhibition of return (IOR, e.g., Rafal et al., 1989), and for this reason could provide less information about the impact of recent trials.

### Estimation of learning rate from Rescorla-Wagner model applied to Fixation Bias data

We evaluated whether the FB data could be accounted for by a Rescorla-Wagner (RW) model, and relatedly, whether a RW-model that reflected a combination of two processes with different learning rates accounts better for the data. The basic model we constructed fit the FB data according to transition probabilities estimated from a RW process, implemented as in Equation 2:

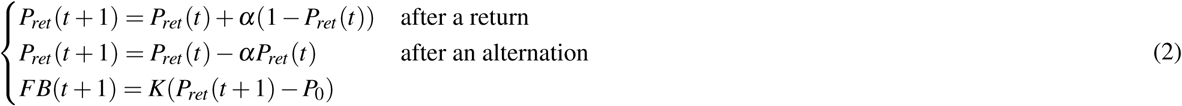

This is a standard RW model, with the exception that it fits anticipatory behavior captured by FB rather than a response to a stimulus. The third line presents the response model that maps the subject’s belief about the transition distribution to the observed fixation biases. it is a simple linear relationship between the internal probability and FB. In Equation. 2, α is the *learning rate, K* is a *scaling factor* transforming internal probability estimates to overt behavior and *P*_0_ is a *probability equilibrium point* reflecting an internal estimate of probability of return above which a participant shows a gaze bias towards the return side. α and *P*_0_ were bounded in the interval [0,1]. We fit the *P*_0_ parameter because it is known that in binary contexts, subjective points of equilibrium significantly deviate from 50%; a truly random binary series is subjectively perceived as having too many streaks (see Falk & Konold, 1997). The reduced model where *P*_0_ was fixed at 50% offered a significantly poorer fit as evaluated by a *Bayes Information Criterion (BIC)* criterion and is not discussed further; Δ*BIC* = 18 ± 5 in pret30 and Δ*BIC* = 16 ± 5 in pret70, both above zero with *p* < .001, bootstrap test.

To evaluate whether FB reflects two learning processes with different learning rates we also fit an extended model in which probabilities were updated based on two processes with different learning rates (see Bornstein & Daw, 2012). In this model, two estimations of the transition probability are updated independently, 
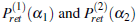
 as in Equation 2, and an overall summary statistic is defined as their weighted average as in Equation 3:

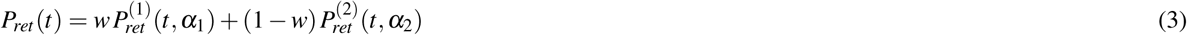

As compared to the simpler model in Equation 2, this model contains two additional parameters; an additional learning rate parameter and a weighting coefficient, *w*. See *Additional Information* for validation procedure details. While the RW model is heuristic in nature, it performs similarly to more complex generative models when the target statistics are stationary (Mengotti et al., 2017).

### Information provided about transition structure by fixation biases and saccade latencies

To evaluate whether FB and saccade latencies provided complementary or independent information about the transition structure in the series, we used a Mutual Information (**MI**) analysis. **MI** captures the amount of knowledge one variable provides about another, or equivalently, the uncertainty about one variable that is reduced by knowing another (Cover & Thomas, 1991). MI does not assume any particular relationship between two variables and captures all orders of correlations, while Pearson’s *R* quantifies a linear relationship (see Equation 4).

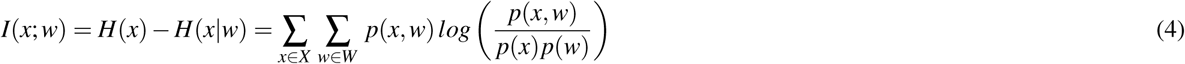

In Equation 4, *H(x)* is the entropy of the variable *x* (here, the experimental condition pret), and *H(x\w)* is the entropy of *x* given *w* (the specific known behavioral response). Because the two stochastic processes (pret70, pret30) were equally probable, the entropy related to which condition participants were observing (*pret* equal to 70 or 30) on any given trial was 1 bit. We used MI to quantify the degree of uncertainty removed about the variable pret by considering several oculomotor information sources and their joint distribution. First we calculated the entropy reduction achieved by FB, *I(pret; FB)*. Second, we performed the same calculation for the saccade latency measure, *I(pret; SL)*. Because saccade latencies on any given trial likely depend on whether the saccade was was an alternate or a return due to IOR, we also partialized by this factor in the MI formulation (see *Additional Information*). Third, we calculated the uncertainty removed when considering the joint (bivariate) distribution of FB and saccade latency, *I(pret; FB&SL)*.

We calculated these three MI quantities per participant, which licensed statistical tests at the group level. We determined. *i*) whether FB and saccade latency were differentially informative with respect to average transition structure and *ii*) whether they provide redundant information (Schneideman, Bialek, &Berry, 2003) about the transition structure, in which case MI provided by the joint distribution is lower than the sum of the two former terms, as shown in Equation 5:

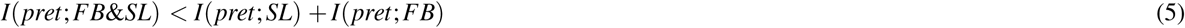

Finally we calculated the information about *pret* carried by separate oculomotor contributions to FB.

### Eye-movement sources underlying FB

This analysis quantified the types of oculomotor movements that may underlie FB. To this end we identified different types of eye movements in the period encompassing the presentation of the fixation symbol and the subsequent pre-target blank screen, and evaluated their direction, using the same coding as FB: positive/negative values for movements made towards/away from the direction of the last target. Here we evaluated whether FB was driven by small involuntary saccadic movements in the range 0.1° – 4.0° observed during fixation (Abadi & Gowen, 2004), as well as small drifts during fixation (Hartmann et al., 2015; Clerici et al., 2002). To avoid contamination of the drift measurement due to the oscillation following the saccade to fixation symbol, we quantified drift only when saccades did not occur. We quantified drift assuming a linear trend; that is, we estimated the initial and terminal eye positions of each drift period via linear regression.

## Extension to four-quadrant design

### Methods

Forty volunteers participated in the study (mean *Age* = 23 ± 5). They were recruited from the local student community, and reimbursed 15 Euro for their time. The Ethical Review Board approved the study.

Participants observed multiple series, each consisting of 128 trials, with the same timing as in Experiment 1. Each trial consisted of a fixation symbol appearing at center, followed by a visual target that appeared at one of four screen quadrants (top left, top right, lower left, lower right), at an eccentricity of 9° from screen center. The target images were unique within each series and belonged to one of four categories. faces, musical instruments, fruits and tools. As in prior work (Davis & Hasson, 2016) we used either highly-constrained or weakly-constrained Markov processes to independently control the predictability of the location-transition and category-transitions presented. Crossing these two factors produced four types of series, where either. *i*) both the location and image category were weakly predictable; *ii*) both the location and image category were highly predictable; *iii*) only location was highly predictable; and *iv*) only category was highly predictable.

Examples of a highly-constraining (high predictability; *HP*) and weakly-constraining transition matrices (low predictability; *LP*) are respectively.

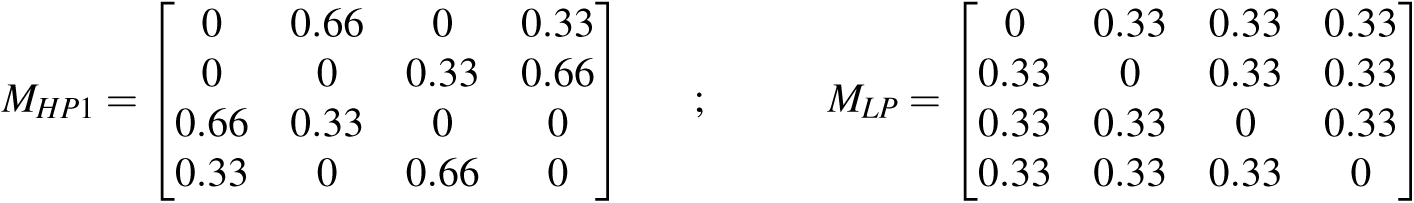

As shown in these matrices, the difference between the two types is that *M_HP1_* contains transitions with 66% probability, and *M_LP_* consists of a uniform set of transitions with 33% probability. Markov entropy was 0.92 bits/trial for the HP process and 1.58 bits/trial for the LP process. In constructing series within each of these 4 conditions, the transitions governing locations and categories were independent (i.e., they were determined by different processes), so that their statistical features needed to be tracked separately. From these transition matrices we produced series with 120 trials, following the same procedure as in Experiment 1, and with the same trial timing. To each 120-trial series, we added 8 trails with random images presented in clockwise or anti-clockwise manner to partially reduce the impact of recent statistical structure. In the study, these series were presented according to a random order determined separately per participant. The first 8 trials of each series were not analyzed as by definition the HP and LP series cannot be discriminated immediately.

Given that our interest is in potential anticipatory biases related to target location, for purposes of addressing gaze data in this study, we ignore the category predictability factor by collapsing across its levels and just examine differences between series depending on whether the location series participants observed was highly predictable (HP) or had low predictability (LP).

The series were constructed so that in both HP and LP, *i*) the marginal frequencies of the 4 locations were 25% (i.e., all four locations were visited equally often independent of the transition patterns), *ii*) returns to the prior location were not possible, and *iii*) The mean frequency of the different pairwise transitions (e.g., a transition from top left to top right location on two successive trials) was equal. We achieved this by creating 4 high-constraint transition matrices so that the distribution of transitions was matched for the high and low predictability conditions (see *Additional Information* for further details). There were 8 series in the HP condition and 8 in the LP condition.

### Analyses

Our first analysis evaluated whether gaze location in the HP condition was strongly impacted by the location of the next most probable target. We coded Gaze Bias as positive/negative depending on whether average gaze location was to the right/left of fixation during the last 10ms of the pre-target blank interval. We analyzed Gaze Bias data using an ANOVA with 3 factors: *i*) vertical position of the next P=66% target (top, down), *ii*) horizontal position of the next P=66% target (left, right) and *iii*) screen-side of just-presented prior image (left, right). We included this last factor because signatures of anticipatory gaze might be weaker when the most probable target location is on the same screen side as the last-presented target, due to IOR effects.

The second analysis used a steady-state approach, where we examined the frequency content of time series of the Gaze Bias in consecutive trials (see *Additional Information* for details). This approach is applicable, because the HP and LP series are generated by different Markov processes and therefore have different recurrence patterns. As detailed in *Additional Information*, the HP and LP target location series have different spectra because for the LP process the modal recurrence time is two trials (the most probable event is to alternate screen side on each trial as two of the three potential targets are on the alternate side); for the HP process, the modal recurrence is 3 (Figure AI.4). If these recurrence patterns drive anticipatory gaze (that is, Gaze Bias), this will directly translate into different power spectra when the anticipatory gaze positions are analyzed in the frequency domain. The steady-state analysis is essentially model-free which means that a stimulus-driven steady-state oculomotor response could be identified even if anticipatory eye gaze patterns are a result of complex computations that cannot be hypothesized in advance. Such complex integration/prediction patterns would rule out any direct relation between anticipatory gaze and the most probable next location, but would still be identifiable in a steady-state analysis. Put differently, even in absence of a specific *a-priori* model of how learning and predictions occur in a complicated non-random context, it would be possible to identify signatures of regularity in the time series of eye data.

## Results

### Fixation Biases reflect stochastic context and structure of recent trials

For all participants, FB was greater in pret70 than in pret30 (Figure 2A. The mean FB difference between the two conditions (Δ*FB* henceforth) was around 0.3° and statistically significant, *t*(20) = 10.10; *p* < .001; *d* = 1.87. In pret70, mean FB was significantly positive, *M* = 0.27 ± 0.04°, indicating a bias towards the side of the last presented target (t-test against zero; *t*(20) = 6.84; *p* < .001; *d* = 1.49). In pret30, the mean FB was negative, *M* = −0.04 ± 0.03°, but not significantly different from zero (*p* > 0.1). Note that despite the statistical significance of the FB effects and robust effect size, the absolute FB values were modest. The average FB was less than 0.4° from center. Thus, FB biases were manifested within spatial zone of the just-removed fixation symbol.^1^

**Figure 2.**
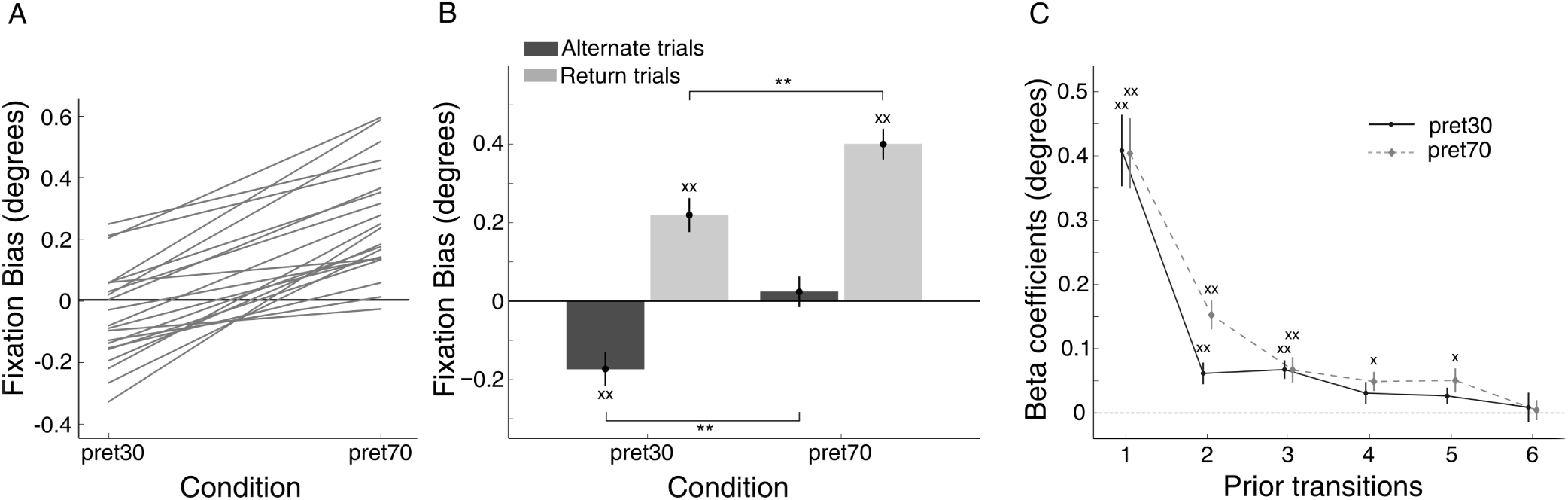
The impact of statistical structure on Fixation Bias. **Panel A:** mean FB values were significantly greater in pret70 than pret30, and the pattern held for all participants (each participant marked via line). **Panel B**: partitioning FB values by most recent transition indicates an effect of statistical structure as well as an impact of most recent transition, as FB was greater immediately after a return than after an alternation. Crosses above each bar indicate significant differences from zero. Asterisks above/below bar pairs indicate significant difference. **Panel C:** A regression model shows that FB was impacted by a return in any of the last 5 transitions for pret70 and in any of the last 3 transitions for pret30.

To evaluate the impact of the immediately prior trial we partitioned FB by the type of prior saccade (return, alternation) and condition (pret70, pret30). Figure 2B presents the mean values for these four cells. A two-way ANOVA with condition (pret70, pret30) and last trial (alternate, repeat) as factors indicated a main effect of condition, as FB values were generally larger in pret70, *F*(1, 20) = 18.1; *p* < .001, and a main effect of last trial as FB values were greater after returns saccades, *F*(1,20) = 75.1; *p* < .001. The interaction term did not approach significance (*F* < 1).

We used regression models to determine the impact of each of the 6 last transitions (i.e., 7 trials) on FB, separately for pret70 and pret30 (see *Methods*). As shown in Figure 2C, in both conditions there was a rapidly decaying impact of recent returns/alternates on current FB, with returns contributing positively to FB. For pret70, the regression model explained 8.0 ± 1.2% of the variance, and the first 5 coefficients exceeded zero (Bonferroni corrected). For pret30 the model explained 7.5 ± 1.3% of the variance, and the first 3 coefficients exceeded zero (Bonferroni corrected).^2^ We additionally evaluated a similar model coding for history of right/left screen sides in recent trials. This model produced statistically significant weights for the two most recent locations, but these effects were an order of magnitude smaller than those found for transition history.

### A Rescorla-Wagner model predicts Fixation Biases

We modeled FB according to a Rescorla-Wagner model, but including two additional parameters: *P*_0_ which reflects the internal subjective point of equilibrium, and *K* which is a multiplicative scaling factor related to transforming internal probabilities to FB magnitudes (see *Methods*). On average, the model accounted for 8 ± 1% of adjusted variance in both conditions. We also compared this model, which contained a single learning rate parameter, to an extended model that reflected a combination of two processes with two learning rates (see *Methods*). Because these models differed in the number of free parameters, we compared their performance using BIC. For both pret70 and pret30, the extended model did not provide a significant improvement in fit (Δ*BIC* not different from zero, p > .05). We therefore report only results for the simpler model as shown in Equation 2.

To evaluate the model, for each participant we used a 10-fold leave-one-series-out validation scheme, where model parameters estimated from 9 series were used to predict FB data in a left-out series. The variance accounted by this procedure exceeded permutation-derived chance (*p* < .05; see *Additional Information*) for 19/20 participants in pret30, and 19/20 participants in pret70 (see Figure 3A for sample prediction of left-out FB data).

Given the validation of the model we then examined the estimated parameters themselves (Figure 3B). The learning rate α was higher in pret30 (*M* = 0.71 ± 0.05) than in pret70 (*M* = 0.55 ± 0.05), *t*(20) = 2.44; *p* < .05; *d* = 0.67. Because we bound the *P_ret_* parameter in the interval [0,1] the range of FB was determined by the scaling factor *K*. We found that *K* was significantly greater in pret70 (*M* = 0.87 ± 0.12) than in pret30 (*M* = 0.49 ± 0.09), *t*(20) = 3.32; *p* < .01; *d* = 0.81. Finally, the mean equilibrium point, *P*_0_, was higher in pret30 (*M* = 0.49 ± 0.06) than in pret70 (*M* = 0.32 ± 0.05), *t*(20) = 2.92; *p* < .01; *d* = 0.69, and differed from 0.5 only for the latter, *t*(20) = 4.06; *p* < .001; *d* = 0.96.

**Figure 3.**
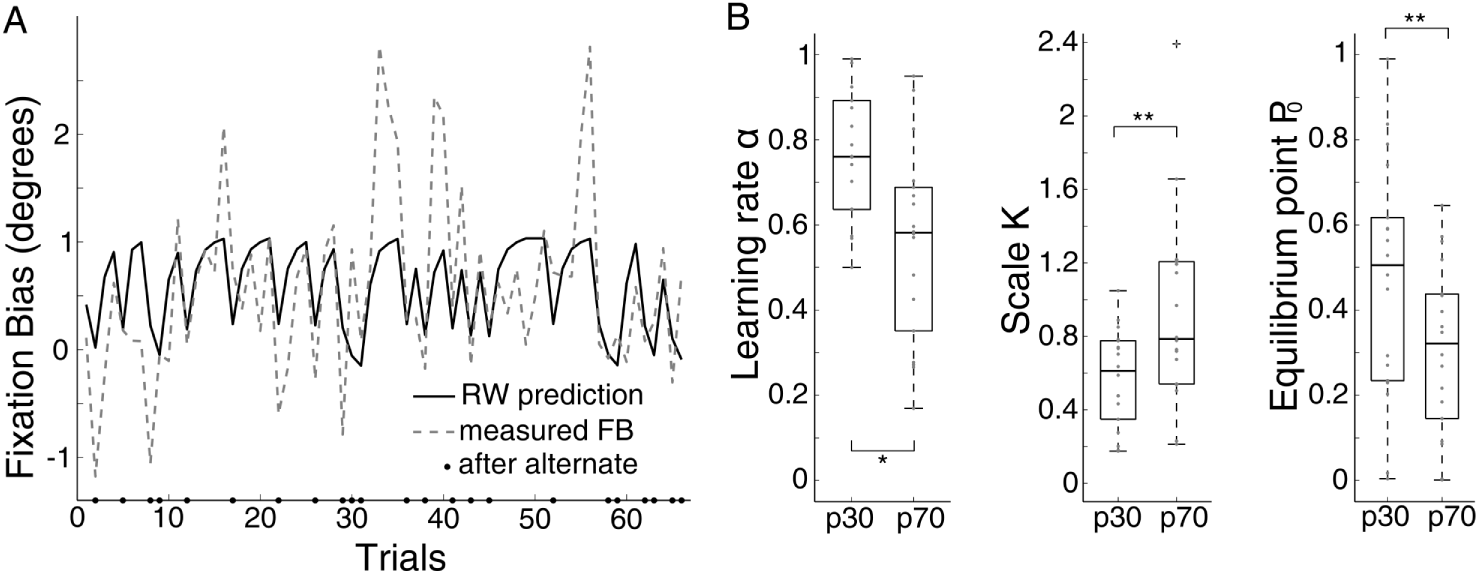
Rescorla-Wagner model of Fixation Bias. **Panel A:** Sample FB data from one session in pret70 condition (dashed line) and model prediction (continuous line) derived from parameters estimated from independent series. Asterisks on abscissa mark alternate (side-switch) trials. Data are concatenated to exclude missing or invalid values. **Panel B**: distributions of model parameters in the two conditions. From the left: learning rate, scaling factor, and equilibrium point. *P*_0_ significantly departed from 0.5 only in pret70. Asterisks above/below bar pairs indicate significant differences.

### Fixation Biases contain signatures of long-term statistical learning

We found evidence for learning of long-term statistical structure in the FB data. We first evaluated FB during the 20 random-location trials that were appended to each series (trials 101–120). Any differences in FB during these trials can only reflect a carry-over effect from the transition structure in the preceding 100 trials. We found a strong carry-over effect, as shown in Figure 4A, with greater FB values following pret70 series. A 2 (Condition: pret30, pret70) x 2 (Last trial. return, alternate) ANOVA confirmed this observation, showing a main effect of condition, *F*(1, 20) = 8.99, *p* < .01. Importantly, this effect was concomitant with an independent effect of last transition in these 20 trials, *F*(1, 20) = 41.31, *p* < .001, because FB was larger after returns. In all, during these random trials, we found a strong effect of the most recent trial, which summed linearly with a longer term impact of the transition structure in the series that preceded the random trials.

Sensitivity to longer-term statistics was also seen in the development of differences between FB values in the pret70 and pret30 conditions (ΔFB) over the 100 trials within each series (see Figure 4B). ΔFB significantly increased from 0.25 ± 0.03° in trials 1–50 to 0.39 ± 0.04° in trials 51–100, *t*(20) = 3.29; *p* < .01, *d* = 0.69. A similar analysis using Mutual Information (see Methods) indicated that FB carried less information about the experimental condition in the first 50 trials (0.029 ± 0.004 bits) than in the last 50 trials (0.054 ± 0.006 bits), *t*(20) = 7.57, *p* < .001, *d* = 2.16 (Figure 4C).

**Figure 4.**
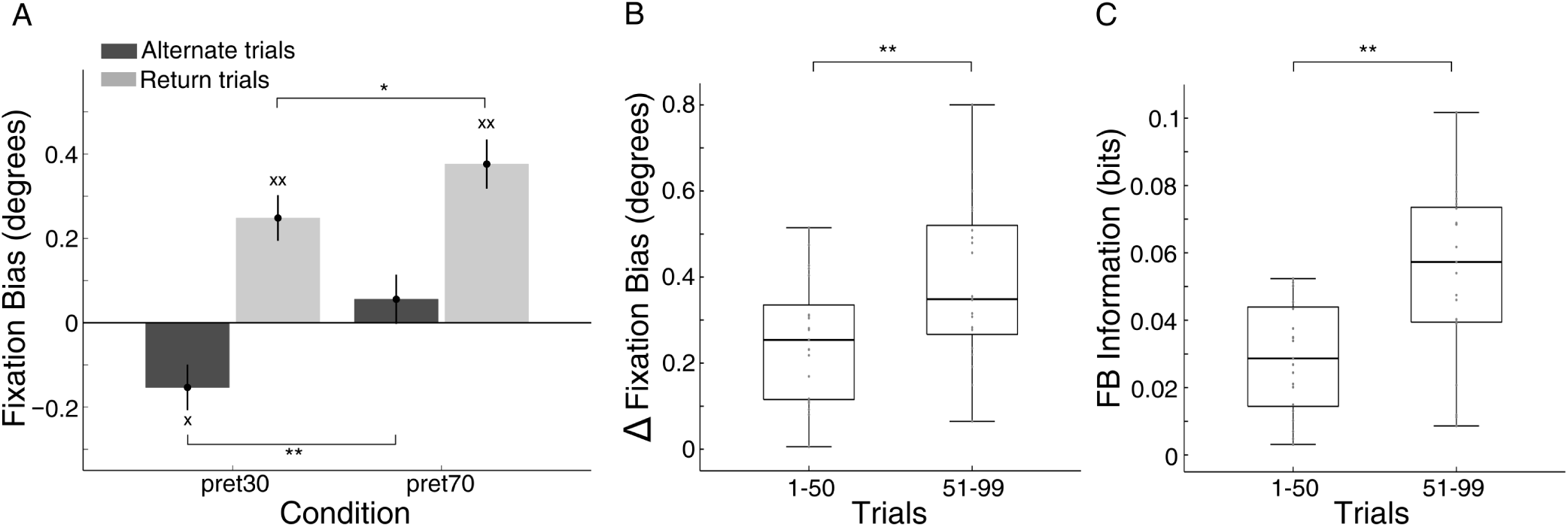
Long term learning signatures in Fixation Bias. **Panel A**: FB values in the 20 random trials (pret=50%) appended to each experimental series. Average FB magnitudes indicate confinement to the area of the fixation symbol (< 0.5° eccentricity). There was a strong impact of the statistical structure of the series presented prior to the random trials, and independently, a strong impact of the immediately preceding trial. Crosses above each bar indicate significant differences from zero. Asterisks above/below bar pairs indicate significant difference (also in following panels). **Panel B**: Δ*FB* was defined as the difference between FB values in the pret70 and pret30 conditions. Its values significantly increased from the first half to the second half of the experimental series. **Panel C**: Similar results when quantified via Mutual Information. In all panels, measures of spread indicate variance within condition and are provided for completeness; they are not indicative of effect sizes in within-participant contrasts.

### Fixation Biases develop within a trial and are co-determined by gaze drifts and saccade instabilities

Before quantifying the development of fixation biases within a trial, we first qualitatively present the trajectories of gaze movements (on the horizontal, *x*-direction), from the point that participants saccaded to center (i.e., time locked to *landing* in the vicinity of the fixation symbol, which tended to occur approximately 10ms in advance of presentation of fixation symbol). Figure 5 presents the time lines of mean gaze location from landing, through the presentation of the fixation symbol and the subsequent blank screen, in 10ms time bins (negative *y* values indicate left screen side, positive values indicate the right side). The process captured by the figure is clear. in both conditions (pret30 and pret70), the landing position (*t* = 0) was on the screen side of the prior target, and in both conditions this was followed by an adjustment towards the screen center during the following ∼ 200*ms*; as we show below these adjustments reflected both drifts and small corrective saccadic movements during the presentation of fixation symbol and the subsequent blank screen. From thereon, gaze trajectories further diverged based on the experimental condition. the gaze stayed on the side of the prior target for pret70, but continued a trajectory towards the alternate side for pret30. For all time points we found a significant difference between the mean Gaze in the two conditions (p < 0.01, Bonferroni corrected). Importantly however, as expressed by the Cohen’s effect size (Figure 5, red line) this difference between conditions showed a continuous increase during the fixation symbol presentation ∼ 400*ms* and during the blank screen ~ 400 – 560*ms*. In *Additional Information* we present density plots of group-level gaze locations in pret70 and pret30, in trials following a target on the left or right screen side (Figure AI.1. It demonstrates the tight clustering of gaze locations at screen center during the window where FB was quantified, as well as the biases induced by transition structure.

We then conducted a quantitative evaluation to determine if the differences in FB between pret70 and pret30 developed between the time of initial landing (FB-landing, prior to presentation of fixation) and the main FB measurement taken during the blank interval after fixation presentation. Finding such a pattern would indicate that any final differences in FB, measured after the disappearance of the fixation symbol, developed from the time of saccade landing. To this end we analyzed the FB data using an ANOVA with three factors, all within-participants. 1) Trial stage: FB measure quantified either after fixation [main FB measure] or at landing; 2) Condition: pret70 or pret30; and 3) Last trial: alternate, return. The ANOVA indicated a main effect of Trial stage, as FB values were generally greater at landing, *F*(1, 20) = 20.8, *p* < .001, reflecting the aforementioned undershoot. There was also a main effect of last trial as bias was greater after return trials, *F*(1, 20) = 23.6, *p* < .001, and a main effect of condition as biases were greater for pret70, *F*(1, 20) = 33.0, *p* < .001. Most importantly, there was also a statistically significant 2-way interaction between Trial stage and Condition, *F*(1,20) = 10.8, *p* = .004, which was due to the fact that the differences in FB values for pret70 and pret30 were smaller when measured at landing than when measured immediately prior to the next target.

**Figure 5.**
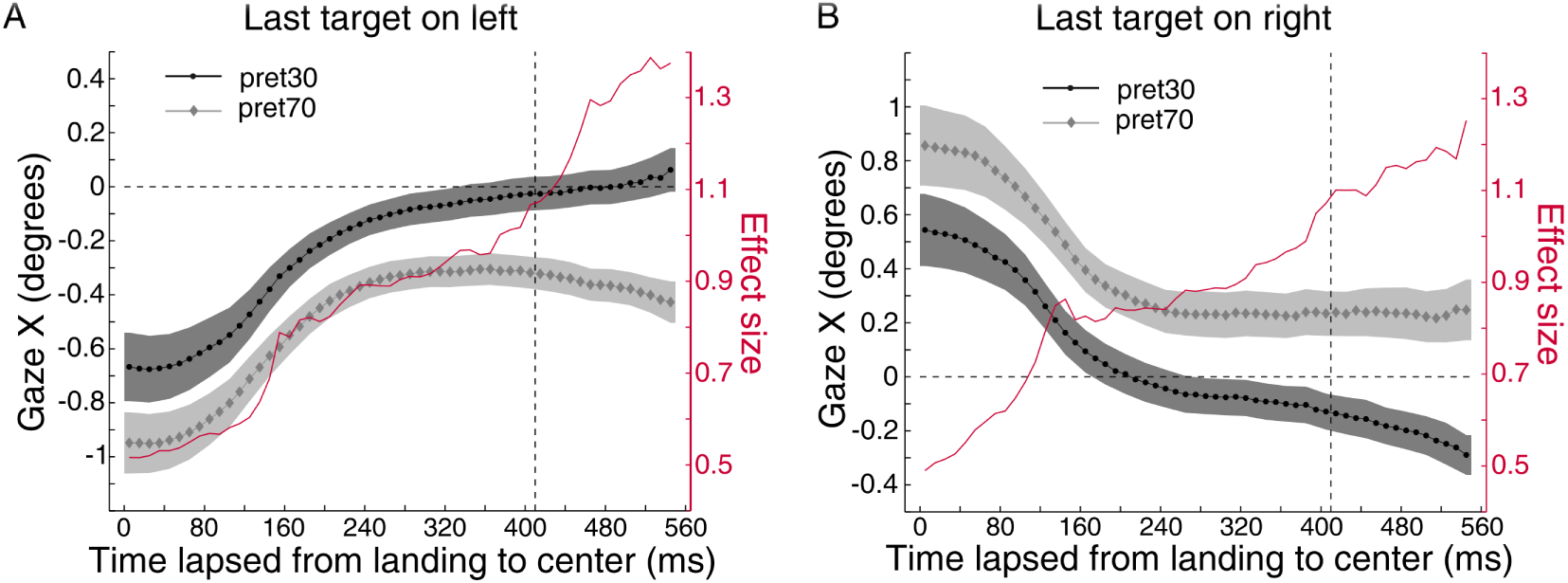
Mean Gaze locked to the time of landing to fixation symbol. **Panel A**: Gaze locations on trials following a target presented on the left. Plots are time-locked to the time at which the saccade to center occurred. Each time point is an average of gaze values in 10ms bins and gaze values occurring during saccades were excluded (shaded area represent ± *SEM*.) Gaze values are coded as positive to the right of screen center and negative to the left. The dashed vertical line indicates the temporal onset of the blank screen (∼ 410*ms* from landing at center). Superimposed (red line; y-axis) is Cohen’s effect size in each time bin. **Panel B**: same as Panel A but for trials following a target on the right.

We also found that, collapsing across condition, the impact of whether the last trial was a return or alternation was weaker when measured at landing (difference ≃ 0.35) than when measured immediately before the next target (difference ≃ 1.1). This produced a statistically-significant two-way interaction between measurement time and Last trial, *F*(1, 20) = 6.6, *p* = .009.

After the gaze arrived at fixation, we observed relatively frequent saccadic intrusions (SI), which are relatively small saccade instabilities in the range 0.1° – 4.0° (Abadi & Gowen, 2004). In both pret30 and pret70, these SIs occurred in a direction opposed to the prior target location. These SI patterns reflected correction to the landing undershoot, seen in that SI magnitudes were strongly negatively correlated with FB landing, indicating more extensive corrections for stronger undershoots (across participants, mean Z-transformed Pearson‘s *R* = −1.1 ± 0.3, significantly below zero, *t*(20) = 17.73, *p* < .001, *d* = 3.87). While the magnitude of SIs was similar across conditions (∼ 1.22° in both conditions), these events occurred significantly more often in the pret30 condition (SI frequency for pret30. M = 1.50 ± 0.08 Hz; for pret70. *M* = 1.39 ± 0.08 *Hz*, *t*(20) = 3.25, *p* < .01, *d* = 0.29). Note that targets were presented at rate of 0.93 Hz. When examining drifts, we found that estimated drift magnitudes were small (< 0.1°), but did differ between pret70 and pret30. A 2 (Condition. pret30, pret70) x 2 (Last trial. return, alternate) ANOVA revealed a main effect of condition, *F*(1, 20) = 9.57, *p* < .01: drifts were negative for pret30 but positive for pret70 (for pret30. *M* = −0.022 ± 0.005°, *t*(20) = −4.13. *p* = .001, *d* = 0.90; for pret70: *M* = 0.011 ± 0.006°, *t*(20) = 1.9, *p* = .05, *d* = 0.43). Separately, drift values were also slightly positive after return trials and slightly negative after alternate trails, resulting in main effect of last trial, *F*(1,20) = 38.8, *p* < .001.

## Saccade latencies

### Averages, LATER model, and impact of recent trials

We concisely report an analysis of saccade latencies (SL) because SL have been used to study the impact of statistical structure and expectation, and because identifying the expected SL patterns would license relating FB to SL data on a trial by trial basis. The statistical structure of the series produced the expected impact on SL (see Figure 6A). A 2 (Condition: pret30, pret70) x 2 (Current trial: alternate, return) ANOVA revealed that, as expected, saccade latencies were slower on return trials, seen in a main effect of current trial, *F*(1, 20) = 14.59, *p* < .001. Speaking to statistical learning, there was also a significant interaction, *F(1,20)* = 6.83, *p* = .01: return saccades were faster in pret70 than pret30, *t*(20) = 5.44, *p* < .001, *d* = 0.48, and conversely alternations were faster in pret30 than pret70, *t(20)* = 5.03, *p* < .001, *d* = 0.72. That is, both return and alternate saccades were performed more quickly in the condition in which they were more frequent.

To understand whether the saccade latencies could be associated with either a threshold shift prior to saccade generation, or accumulation rate, we fit a LATER model (Carpenter & Williams, 1995) to SL data in each of these four conditions, solving for threshold (*ϑ*) and accumulation rate (*μ*) (see *Additional Information*). A 2 (Condition: pret30, pret70) x 2 (Current Trial: alternate, return) ANOVA on the estimated threshold parameters *ϑ* revealed a significant two-way interaction, *F(1,20)* = 14.37, *p* < .001, because the threshold parameter strongly tracked stimulus likelihood, but was not sensitive to whether the saccade was an alternate or return. Specifically, in pret30, thresholds were significantly lower for alternate saccades than returns (difference = 0.10 ± 0.05, *t(20)* = 2.13, *p* < .05, *d* = 0.90). Conversely, in pret70, thresholds were significantly higher for alternate saccades than returns (difference = 0.06 ± 0.03, *t(20)* = 1.98, *p* < .05, *d* = 0.78). A very different pattern was found for accumulation rate (*μ*), where the ANOVA identified only a main effect of current trial (return vs. alternation), *F(1,20)* = 7.79, *p* < .01, indicating more rapid accumulation for alternate saccades. Here, there was no significant interaction.

We used regression models to determine the impact of recent transitions on SL (see Figure 6B). Given the strong impact of IOR on SL, we fit separate regression models to alternate trials and return trials in pret70 and pret30. IOR could reduce the sensitivity for identifying recent-trial effects when analyzing SL for return saccades. For return saccades in pret30, only the lag1 coefficient significantly departed from zero, because return saccades were faster when preceded by a return, *t*(20) = 4.04, *p* < .01, *d* = 0.88. The model for alternation saccades in pret30 did not reveal any impact of prior trials. Note that the same model applied to FB data in pret30 indicated sensitivity to the last three transitions. For alternate trials in pret70 the coefficients from lag-1 to lag-4 were significantly positive indicating that alternation saccades were slowed down by a return saccade in any of the four prior transitions.^3^ This was roughly comparable to the findings for FB, which indicated sensitivity to the last 5 prior transitions in this condition. The model fit for return saccades in this condition did not reveal any impact of prior trials. In summary, for saccade latencies evidence for the impact of prior trials was only found for transitions that were less expected in a given condition (returns in pret30, alternates in pret70).

**Figure 6.**
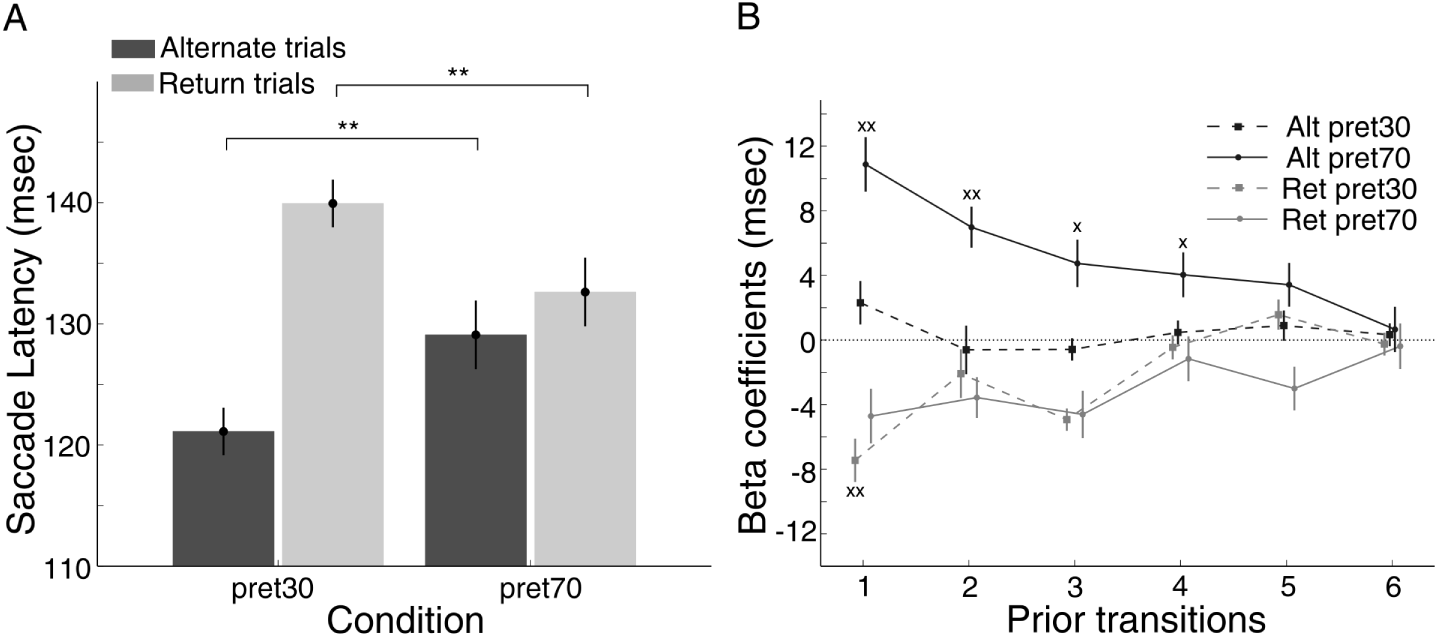
The impact of statistical structure on saccade latency. **Panel A**: Saccade latencies indicate learning of statistical structure in addition to an effect of whether a saccade is a return or alternation. Asterisks above bar pairs indicate significant difference. **Panel B**: Regression models for saccade latencies in the pret30 and pret70 conditions, constructed separately for alternate (Alt) and return (Ret) trials. For pret70, a return in any of the last 4 transitions impacted positively (slowed down) SL in alternate trials. For pret30, return saccades were faster when preceded by a return saccade in the immediately prior trial, but there was no indication for more remote effects. Crosses above each point indicate significant differences from zero.

### Signatures of learning long-term statistics in saccade latency

Examining saccade latencies to alternations and returns during the 20 random trials appended to each series, we found that sacccades on alternation trials were faster after the pret30 condition than after the pret70 condition, consistent with a carry-over effect, *t*(20) = 2.45, *p* < .05, *d* = 0.45. However saccades on return trials were not faster after pret70 than after pret30.

Second, we evaluated whether indexes of learning developed from the first half of each series (trials 1-50) to the second half (trials 51-100). We defined the following contrast term as an index of statistical learning.

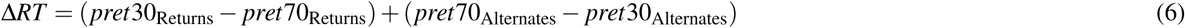

ΔRT is larger to the extent that individuals can compute what is the more probable transition in each condition. Consistent with statistical learning, ΔRT significantly increased from the first to the second half of the 100-trial sessions (first half = 14 ± 3*ms*; second half = 20 ± 3*ms*; *t*(20) = 6.45, *p* < .001, *d* = 0.43). In a similar analysis we quantified the Mutual Information between SL and the experimental condition, conditioned on whether each trial was as return or alternations: *I(pret; SL|S)*, where returns are coded as S=1 (see *Additional Information*). Mutual information significantly increased from the first 50 trials, *I(pret; SL|S)* = 0.012 ± 0.005 *bits*, to the last 50 trials, I(pret; SL|S) = 0.016 ± 0.006 *bits*, *t*(20) = 2.60, *p* < .01, *d* = 0.82. This confirms that saccade latencies provided more information about the statistical structure of the series in the second half of the trials.

Nevertheless, when considering these latter MI values in relation to the ones we found for the FB data, it is notable that for trials 51–100, fixation biases provided around three times the information about the statistical process, 0.054 bits for FB vs. 0.016 bits for SL, *t*(20) = 6.27, *p* < .001, *d* = 1.50.

### Trial-level correlations between Fixation Bias and saccade latency

We evaluated the correlation between trial-by-trial FB values and the latency of the immediately subsequent saccade. Recall we defined FB as a bias towards the side of prior target. If FB is related to anticipation of a screen side, then greater FB values should be linked to *1*) faster latencies for return saccades and conversely, to *2*) slower latencies for alternate saccades; this should hold for both pret70 and pret30, to the extent that trial-level fluctuations reflect local preferences for returns or alternations. We evaluated these correlations in the pret30 and pret70 conditions.

The analysis produced two findings (see Figure 7). For pret70, the correlations exactly matched the predicted relation between FB and SL: increased FB was associated with faster latencies for return saccades (across participants, mean z-transformed Pearson‘s *R* = −0.05 ± 0.02, *t*(20) = 2.10, *p* < .05, *d* = 0.46), and slower latencies for alternate saccades (across participants, mean z-transformed Pearson‘s *R* = −0.13 ± 0.02, *t*(20) = 5.20, *p* < .001, *d* = 1.13). For pret30, increased FB was associated with faster latencies for return saccades (across participants, mean z-transformed Pearson‘s *R* = −0.06 ± 0.02, *t*(20) = 2.69, *p* < .01, *d* = 0.58). However, increased FB was not associated with slower saccade latencies to alternate trials. Reflecting these significant trial-by-trial correlations, FB and SL conveyed redundant information so that, *I(pret; FB&SL)* < *I(pret; FB)* + *I(pret; SL|S)*), *t*(20) = 7.19, *p* < .001, *d* = 1.57. The degree of redundancy was moderate however (∼ 20% of *I(pret; FB))*, suggesting SL and FB may convey substantially different information about the target location statistic.

**Figure 7.**
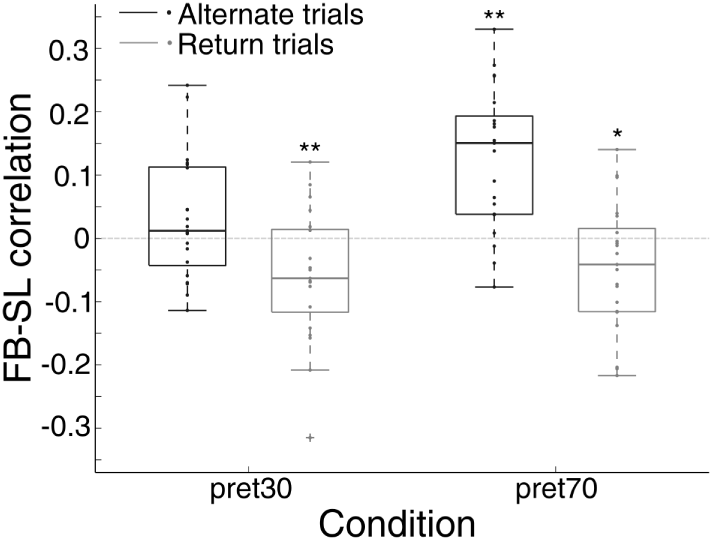
Trial-level correlations between Fixation Bias and saccade latency. Distributions are plotted for pret70 and pre30, partitioned according to whether the saccades were return or alternate saccades. Asterisks above bars mark significant difference from 0 which held in all conditions apart from alternate trials in pret30.

While the trial level correlations between FB and SL held as described above, these relations did not hold when evaluated from an inter-individual perspective. Across participants, increased sensitivity to statistical structure, as measured by ΔFB did not significantly correlate with increased sensitivity to statistical structure as measured by ΔRT. Similarly we did not find significant correlations between the information-theoretic measures *I(pret; FB)* and *I(pret; SL|S)*.

## Extension to four-quadrant study

### Gaze Bias does not directly signal screen side of most likely subsequent target

In the high predictability condition (*HP*), on each trial there was a probability of 66% of transitioning to one location, 33% probability of transitioning to another, and 0% transition to a third (in addition, repeats were never allowed). To understand if anticipatory fixation biases were aligned with the location of the most probably future target, we partitioned all the trials into 4 bins, depending on the most probable target location in the next trial (*P* = 66%). If participants’ gaze tracked the most likely future position then, at first approximation, gaze location should show greater bias towards the right side when the *P* = 66% transition is expected to be on the right than when the *P* = 66% transition is expected to be the left, and similarly, show greater bias towards the top/bottom of the screen depending of the expected vertical position of of next most probable target.

We coded Gaze Bias X during the blank screen prior to target presentation as positive if to the right of fixation and negative if to the left of fixation. An Analysis of Variance with three factors (horizontal position of next most probable target, vertical position of next most probable target, and screen side of just presented image) did not produce any result suggesting that anticipatory gaze was biased towards the horizontal position of the next most likely target. Nor did this effect interact with the other factors. The only significant effect was the location of previous screen, *F*(1,39) = 80.4 *p* < 10^−6^, which indicated a shift towards the side opposite to that of the last presented target, and could reflect a general prediction of screen side alternation.^4^ Figure AI.2 presents Gaze Bias probability densities according to the location of the most likely next target, and does not reveal any observable deviation from the screen center.

### Steady-state analysis identifies tracking of statistical structure

The above analysis did not identify what might be the most straightforward signature of anticipation. gaze did *not* signal the location of the next most probable target. However, as demonstrated in Experiment 1, gaze biases are sensitive to prior trial history (consistent with Bornstein & Daw 2012; Yu & Cohen, 2008). They may also reflect weighted averaging over the set of potential transitions. All this precludes a simple model where gaze location directly tracks the location of the next most predictable target. To determine whether anticipatory gaze biases tracked the transition structure of the experimental series we examined whether these gaze biases tracked the different recurrence features of the Markov processes producing the HP and LP series. We employed an analysis technique that quantifies steady-state responses in a system of interest. This analysis identifies signatures of frequency characteristics in a time series, which are linked to the recurrence rate of an external stimulus. Here, recurrence rate is related to the mean number of trials between repeated presentation of a stimulus at the same location (see *Methods*).

We calculated the power spectral densities of Gaze blank *X* (Gaze blank *Y*) in HP and LP and we defined Δ*PSDx* as the difference between PSDx in HP and LP (and similarly defined Δ*PSDy*, see *Additional Information*). Figure 8 shows the relevant PSD plots for the X and Y gaze dimensions. As shown in the Figure, we found statistically significant differences between the spectral features of the gaze locations in HP and LP. For both PSDx and PSDy we found a significantly stronger peak at a frequency of 1/3 trials for HP than for LP. This is precisely the frequency that characterizes the target-location series in HP (for *X*:*t*(39) = 2.99, *p* < .01, *d* = 0.38; for *Y*:*t*(39) = 2.48, *p* < .05, *d* = 0.31) (see also Figure AI.4). This indicates a differentiation between HP and LP in the frequency representation of gaze positions across trials.

**Figure 8.**
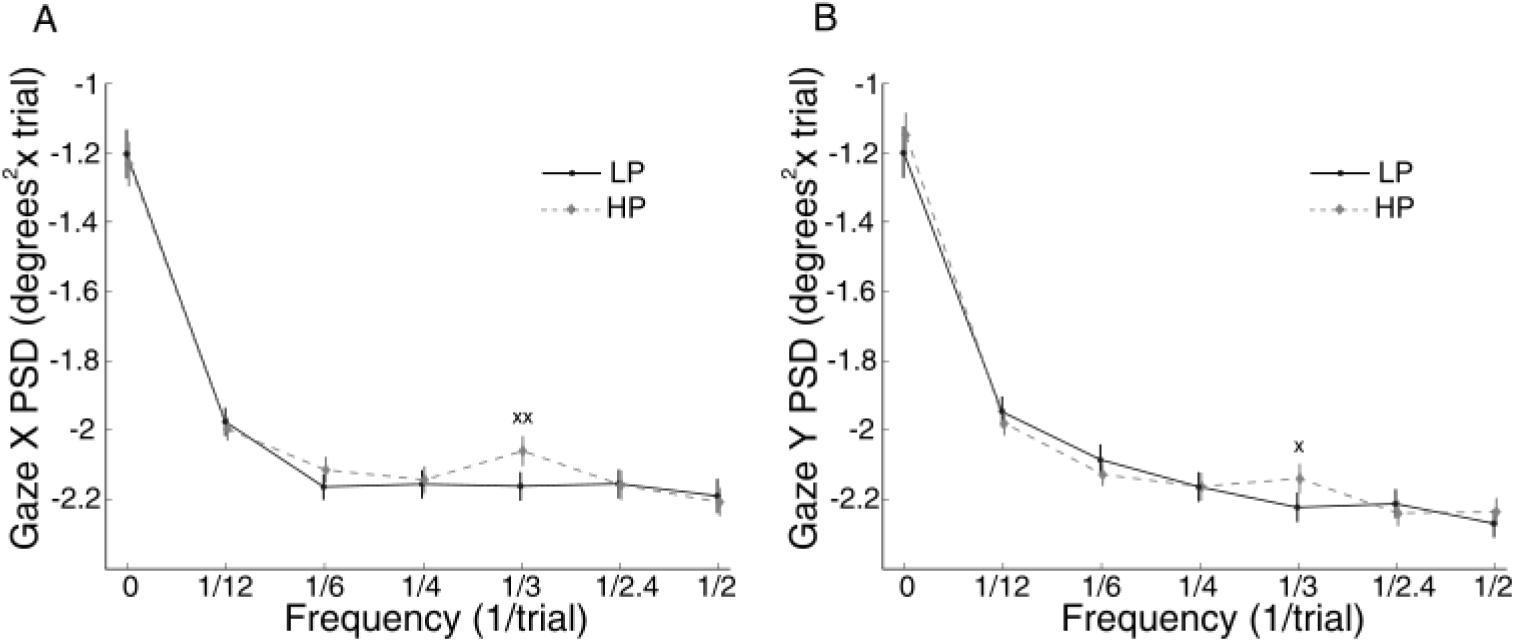
Power Spectral Density of gaze-location series in the High Predictability (HP) and Low Predictability (LP) location series. The analysis was applied separately to the X and Y values of gaze locations measured during the blank screen prior to each target. A single {x,y} tuple was recorded per trial. Crosses above each point indicate significant differences between conditions.

We also applied this steady-state analysis separately for the first and second halves of each series. We found no differences between the HP and LP processes in the first half. However, in the second half of trials Δ*PSDx* at *freq* = 1/3 was significantly greater than zero, *t*(39) = 2.98, *p* < .01, *d* = 0.50. Δ*PSDy* at this frequency was also significantly greater than zero, *t*(39) = 2.07, *p* < .05, *d* = 0.36.

As Figure 8 shows, the power spectra of the gaze-location series reflect the different cross-trial trajectories of anticipatory oculomotor responses in HP and LP. To directly link these gaze positions to the series of target locations in the the two conditions, we performed a coherence analysis to quantify the impact of target locations on Gaze Bias, assuming a linear relationship. We first derived the coherence between the target x-position and Gaze Bias X (*coh_Gaze X*) as well as the coherence between the target y-position and Gaze Bias Y (*coh_Gaze Y*). As a summary statistic we defined total coherence (*tot_coh*) as *coh_GazeX* + *coh_GazeY* (see *Additional Information*). When calculated for the entire series, we did not identify significant difference in the coherence between target and gaze location for HP vs. LP. However, splitting the trials into 1st and 2nd halves, we found a difference in tot coh between HP and LP in the 2nd half, at the *freq* = 1/3 trials, *t*(39) = 3.13, *p* < .01, *d* = 0.49. This shows that the target sequence presented on the screen were driving Gaze Bias in HP process, and this was identifiable during the second half of each series.

## Discussion

There exists extensive literature on how learning impacts response components (Kim et al., 2017; O’Reilly et al., 2013; Vossel et al., 2014) and the brain regions that mediate these responses (e..g Mengotti et al, 2017). Despite these advances, and demonstrations that strong predictability can produce anticipatory motor behaviors (e.g., Dale et al., 2012; Vakil et al., 2017) the impact of learning on predictive processes *per se* remains an open question. This may be due to difficulty in isolating overt behaviors that are informative of predictions, but de-coupled from stimulus responses. Consequently, current theorizing is largely informed by analyses of behavioral or neurobiological *responses* to stimuli that vary in predictability (e.g., den Ouden et al., 2010; Kim et al., 2017; Vossel et al., 2014). Our findings, based on a new oculomotor metric (FB), directly address three core issues on the interface of learning and prediction: i) the prevalence of predictions: transition probabilities strongly impacted FB, and the difference across conditions (ΔFB) had high split-half reliability; ii) the temporal integration-constants of learning: FB contained information about learning on two temporal scales: a micro-scale encompassing events in the recent 4–6 trials, and a macro-scale reflecting features of the stationary distribution from which trials are drawn; and iii) the information carried by predictive vs. stimulus-linked behavior: FB carried significantly more information about the environment’s transition structure than SL, though the latter was also sensitive to transition structure, and there was a trial-by-trial correlation between FB and SL in three of the four examined cases (see Figure 7).

All these findings were obtained via analyses of very subtle fixation biases that were recorded while anticipating targets, with magnitudes of around 1 degree, in a design where predictions could only be based on transition structure.

### Relation to prior work on saccade latencies

There is considerable interest in the perceptual inferences underlying saccades and the temporal time scales governing them. Noorani & Carpenter (2016) review a set of factors that impact SL, which include expectation, urgency and stimulus features. Consistent with several studies (Vossel et al., 2014; Kim et al., 2017; Farrell et al., 2010), we found that saccade latencies indicated learning of statistical structures. return saccades were faster in pret70 than in pret30, and conversely, alternate saccades were faster in pret30 than in pret70. These findings are consistent with several studies. Kim et al. (2017) showed that prior probability for a particular left or right saccade was directly reflected in the rise-to-threshold parameter of a LATER model. Farrell et al. (2010) manipulated the probability of return to the same location in a saccade sequencing paradigm, and modeled SL with a competitive race-to-threshold model (Brown & Heathcote, 2008). They found that the threshold changed with a target’s probability of return and that accumulation rates decreased for return saccades. These findings are very similar to those we obtained when applying a LATER model, which identified the same dissociation. thresholds were reduced by expectation whereas accumulation rate was impacted by whether the saccade was a return or not. In all, the analysis of saccade latencies dovetails with recent conclusions about factors that impact threshold and accumulation rates in non-random environments.

In addition, we found that the structure of recent trials impacted SL. This was most strongly exhibited for alternate saccades in the pret70 condition, which were impacted by return saccades in each of the last 4 transitions. Interestingly, similar trial-history effects were not found for (the more frequent) return saccades in the same condition; documenting these effects for alternate but not return saccades indicates that, in general, the pret70 condition was definitely associated with cumulative integration of recent past trials, but that the behavioral expression of this integration varied as a function of the behavioral response. Saccade latencies also signaled learning on a longer temporal scale: regularities produced stronger signatures of statistical learning (ΔRT) in the second half of each series than in the first, and there was a carryover effect from the statistical structure of the pret70 and pret30 series to the 20 random trials (washout period) that followed those series.

Taken together, our findings indicate that, with respect to established saccade latency measures, our paradigm produced data consistent with learning of transition probabilities between screen sides. We emphasize that in this paradigm, the exact target location could never be predicted because targets were positioned randomly on a 20° arc. Thus, the findings derive from general learning of screen-side transitions rather than the learning of saccade sequences to specific screen locations.

The neural sources that mediate the impact of learning and expectation on saccade execution have been examined in several studies using cue-target or smooth pursuit paradigms, in both humans and non-human primates. Primate studies show that anticipatory signals can be observed in multiple brain systems involved in target selection. The discharge of neurons in superior colliculus (SC) tracks the intended gaze movement (Hafed et al., 2008) and can predict the direction of a cued target (Horwitz & Newsome, 2001). This activity increases with the probability of target presentation in the neuron’s receptive field (Basso & Wurtz, 1998) and is also found during anticipatory periods prior to target appearance (Dorris & Munoz, 1998). Furthermore, during smooth pursuit, some SC neurons increase their firing rate if their receptive field is in the expected location of the next saccade (Dash et al., 2016). Other brain areas have been shown to code for expected locations prior to target appearance (caudate nucleus in Lauwereyns et al., 2002; LIP in Shadlen & Newsome, 2001), likely increasing the pre-target baseline activity of eye effectors (Dorris et al., 1999; Noorani & Capenter, 2016).

That said, stimulus probability can impact post-stimulus responses without being accompanied by signatures of anticipatory activity. In one study (Stadler et al., 2016), the ERP P300 evoked potential linearly tracked stimulus probability, but pre-stimulus activity indexed by the Contingent Negative Variation component did not, but only signaled whether probability differs from zero. Cashdollar et al. (2016) identified robust MEG signatures that differentiated responses to stimuli presented in random vs. regular series, but the pre-stimulus patterns in those series were less differentiated, and mediated by working memory capacity.

Identifying anticipatory behaviors linked to stimulus probability would be useful for separating contributions linked to anticipation from those linked to stimulus response, as well as for identifying neural systems that optimize oculomotor function in predictable contexts. We thus shift to our research focus on Fixation Bias.

### Fixation Biases. inter-trial effects and learning

In terms of absolute magnitude, Fixation Biases were subtle, with 90% of all gazes falling in an area of ±1.6 degrees from center. FB significantly differed between pret70 and pret30, with pret70 linked to a stronger bias towards the screen side of last presented target. FB was also strongly impacted by the most recent trial. returns induced a significant FB towards the last screen side, but more strongly for pret70. Regression models indicated longer-term effects, pointing to independent effects of each of the last five transitions for pret70 and each of the last three transitions for pret30. These are consistent with analyses of behavioral and neurobiological response patterns documented in studies of model-free reinforcement learning (Bornstein & Daw, 2012; Harrison et al., 2011) which showed a rapidly decreasing effect of recent trials.

We also found signatures of learning over longer scales. Differences between FB for pret70 and pret30 (ΔFB) were larger when computed from trials 51–100 than from trials 1–50, a pattern consistent with a conceptually similar analysis we conducted for SL. Furthermore, during the 20 random trials appended to each series, FB was impacted by the preceding statistical structure. Specifically, when the random trials were appended to the pret70 series there was still greater bias towards the last screen location, and when they were appended to the pret30 series, there was still a greater bias towards the alternate side. During the random trials, this continuing long-term impact of the prior statistical structure coexisted with a second, independent effect of whether the last trial was an alternate or return. This indicates that the impact of prior statistical structure, which at that point was not reinforced but memory enabled, maintained above and beyond an independent strong modulation of each prior trial.

While this study constitutes an initial examination of anticipatory FB and its implications for models of learning and prediction, the data produced findings that bear on the relation between formal uncertainty, subjective uncertainty and prediction. Formally, the two Markov processes used here, pret70 and pret30, have equal uncertainty. That is, they have the same marginal frequencies (which could be quantified via Shannon’s Entropy, here maximal at 1 bit) and the same first-order Markov Entropy. Notwithstanding these formal equivalences, pret30 and pret70 were associated with substantially different learning characteristics. Decades of research have repeatedly shown that humans show a specific bias in their judgments of randomness for binary series such as the ones used here: they judge random series as overly regular in that they misperceive them as containing more streaks (repetitions) than one expects from chance (e.g., Falk & Konold, 1997; Williams & Griffiths, 2013). Conversely, humans judge binary series as random only once they contain 60 – 70% alternations. If such biases are not limited to judgments or reasoning, but also impact online learning, then the pret70 and pret30 should be associated with different learning trajectories, with the latter reflecting signatures of a (subjectively) random process.

This was exactly what we found. Fixation biases in pret30 and pret70 were associated with different learning characteristics, in a manner consistent with the aforementioned studies on judgments of randomness. We applied a Rescorla-Wagner model to FB data, which was successfully validated on out-of-sample data for almost all participants. The parameter fits indicated a significantly higher learning rate α for pret30 than pret70, reflecting a narrower temporal integration window in pret30. This result was consistent with the regression model results for FB and SL, which showed a weaker impact of recent trials in pret30. Additionally, the parameter *K*, which reflects the transformation from subjective probability to FB, was significantly larger for pret70 than pret30. This means that, all else being equal, the transformation from the subjective probability estimate to anticipatory behavior was associated with larger scaling effect in pret70. It remains to be determined whether these findings for *K* reflect different levels of confidence in the internal distributional estimations (as captured, e.g., by hyperparameters in Dirichlet distributions), or a difference in how distributional information translates into oculomotor commands. Finally, the findings for the equilibrium point *P*_0_ only partially confirmed expectation. Because prior work suggests that series are perceived as random when the proportion of returns is around 30%, we expected *P*_0_ to be in that range for both conditions. While *P*_0_ differed between the conditions, the distribution in pret30 was qualitatively larger (encompassing almost the entire [0,1] interval, and more work is needed in order to determine this issue.

With respect to potential for future discovery, we showed that analyses of gaze biases can determine whether environmental regularities impact anticipatory behavior, even in absence of an a-priori learning model, or a model of how learning impacts anticipatory behavior. In an extension to the main study, we presented targets in one of four quadrants based on a first-order Markov process. Even though the series of target locations was stochastic rather than deterministic, gaze locations tracked the recurrence characteristics of the Markov process, as reflected in the cyclical nature of target locations. We identified this using a steady-state analysis, which indicated that the gaze-location time series had higher power in the recurrence frequency of the high probability process. Furthermore, we directly linked between the sequence of target locations and the sequence of gaze locations by confirming their coherence in the frequency domain. In contrast, a more conventional analysis, which assumed that gaze bias would be strongly determined by the screen side of the next most probable transition (implemented via an ANOVA), failed to account for significant variance. This suggests that gaze biases in such contexts are not random, but reflect relatively complex integration dynamics.

### Fixation Biases. intra-trial dynamics and relation to oculomotor bases

Fixation Biases were the end-state of oculomotor processes that took place between the offset of one target and the onset of the other. The saccades to center occurred on average slightly (∼ 10*ms*) prior to the appearance of the fixation symbol, indicative of saccade planning that began around 60–70ms prior to fixation onset. The landing saccades undershot fixation by about 1°, with a larger undershoot after return than alternate saccades. Our analyses indicated that the difference in gaze patterns and fixation biases in the pret70 and pret30 developed over the subsequent 560ms (i.e., the combined period of the fixation screen and 160ms of blank screen) culminating in the significant difference documented in the main analysis. Interestingly, the impact of the last saccade on FB (return vs. alternation) also developed over the course of the trial. It was weaker when measured at landing prior to presentation of the fixation cross than when measured during the last 10msec of the pre-target blank screen. These findings suggest that, as compared to the final gaze position (where FB was measured), the initial gazes made to center were more weakly impacted by both the general stochastic context and the type of last trial. These effects developed over the duration of the fixation symbol and subsequent blank screen.

We identified two sorts of oculomotor movements, both of which were impacted by statistical regularities. First, in pret30, there was a greater frequency of small corrective saccades away from the location of the prior target. Second, in pret30 the direction of eye drifts was in the direction opposite to that of the last target, whereas in pret70 drifts were in the same direction.

Drifts and small saccadic instabilities are prevalent during fixation (Cherici et al., 2012), are thought to provide optimal retinal input for downstream visual processing (Rucci & Victor, 2012) and maintain a balance between fixation and anticipation (Watamaniuk et al., 2017). Covert attentional shifts and the execution of eye movements are thought to share several functional networks (Corbetta et al., 1998; Nobre et al., 2000). Micro-saccades are known to associate with anticipated target location in attentional cue-target paradigms (e.g., Meyberg et al., 2017), consistent with our findings. A potential locus of brain activity that underlies FB may be the deep Superius Colliculus, since local pharmacological interventions in primates can produce departures from fixation target (Goffart et al., 2012). However, to our knowledge neural activity in this area has not been correlated yet with oculomotor fixation adjustments.

### Fixation biases and saccade latencies. two sides of the same coin?

The stochastic context impacted both fixation biases and saccade latencies but with notable differences. First, when examining trials in the second halves of the series, for which the impact of experimental condition were more robust, we found that FB conveyed three times more information about the experimental condition (pret30, pret70) than did saccade latencies (the latter conditionalized on whether the saccades were returns or alternates). This difference was consistent with several data patterns suggesting that FB was more sensitive to recent events. The regression models showed that compared to SL, FB was impacted by a more extended trial history in both conditions, consistent with a longer integration window.

The trial-by-trial correlations between FB and SL were consistent with the idea that FB signals an anticipatory prediction, with higher FB values preceding both faster return saccades and slower alternation saccades. That said, the correlation magnitudes were modest, though significant on the group level in most cases. Overall SL and FB had relatively little redundancy with respect to the information they contained about the experimental conditions (∼ 20% of the information carried by FB), suggesting moderate complementarity. Assuming that FB is a signature of information accumulation and prediction, this also suggests that the relationship between FB and SL is complex, possibly reflecting nonlinearities.

## Summary

Our studies show that a proactive oculomotor metric, quantified via subtle anticipatory fixation biases, is strongly impacted by input statistics. These biases were on average less than 1° in magnitude, were measured while participants were fixating the screen center, and presented strong split-half reliability. While fixation biases were moderately predictive of subsequent saccade latencies on a trial-by-trial level, they captured more information about input statistics than did saccade latencies. Finally, trial-level fixation biases contained information about recurrence features of a more complex generating processes, which could be quantified in absence of an explicit a-priori learning model. These results show that strictly anticipatory behavior is impacted by learning on multiple scales, and that fixation biases offer a unique and sensitive avenue for understanding learning and prediction in a way that is decoupled from stimulus-response.

## Additional Information

### Group-level gaze-location density maps

Figure AI.1 presents gaze location patterns during the pre-target blank interval in which we measured FB in Experiment 1. To sample Gaze Bias in an adequate spatial resolution (0.1 × 0.1 degrees^2^), we collapsed data across participants (32752 points). For each condition, we partitioned the fixation data based on screen-side of prior target. The figure communicates that: *i*) the area with maximal density was always at center (0, 0), demonstrating participants’ success in maintaining fixation near the center of the fixation symbol, *ii*) Gaze Bias density steeply decreased in surrounding areas to 1/5 of maximum density, and *iii*) mean Gaze Bias (red crosses) was qualitatively shifted in the direction of the most likely next target location. Figure AI.2 was derived with same procedure from data collected in the 4-quadrant study (44128 points), partitioning Gaze Bias values according to the most likely next target location in the HP condition. There was a relatively smooth decrease in density from screen center to surrounding areas (1/2 of maximum) but no visible shift of mean Gaze Bias (red crosses) in the direction of the most predictable target location.

**Figure AI.1.**
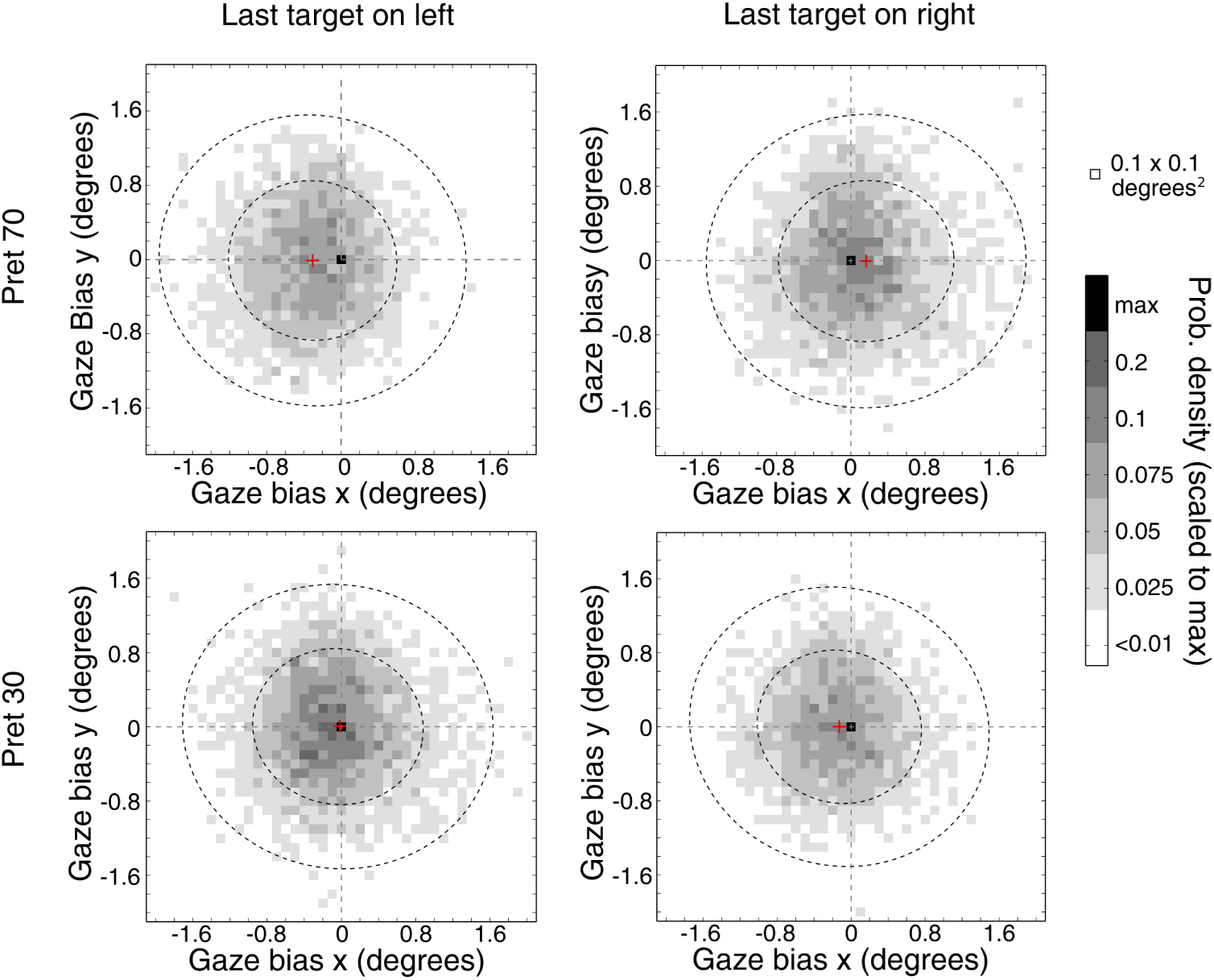
Fixation location during last 10ms of pre-target blank screen in Experiment 1. To present the effect of stochastic context, fixation locations are presented as function of prior target location. Densities were calculated in 0.1 × 0.1 degrees^2^, merging data points from all participants and normalizing to the maximum value for condition. The single dark point marks maximal density and is always at the screen center; red crosses indicate mean values for condition; inner/outer circles mark areas encompassing 50% and 90% of all fixations. In pret70, gaze locations are slightly, but notably shifted toward the side of the last presented target.

### Impact of eccentricity criteria on ΔFB

In the main analysis, we considered trials as valid for FB analysis if the gaze was within 3° from center. To evaluate whether the findings generalized beyond this criterion, we also examined only *i*) the set of trials where the gaze was less than 1.2° (i.e., the eye location was within the area of the just-presented fixation symbol; analyzed trials: 59 ± 3%) and *ii*) the set of trials where gaze was within 5° of fixation (analyzed trials: 66 ± 3%). In all cases we found that the resulting ΔFB was significantly above zero, indicating an impact of statistical structure on FB (see Figure AI.3). In all cases ΔFB significantly increased (*ps* < 0.01) from the first half of the series (trials 1–50) to the second half of the series (trials 51–100), indicating learning over time, as documented in the main analysis.

**Figure AI.2.**
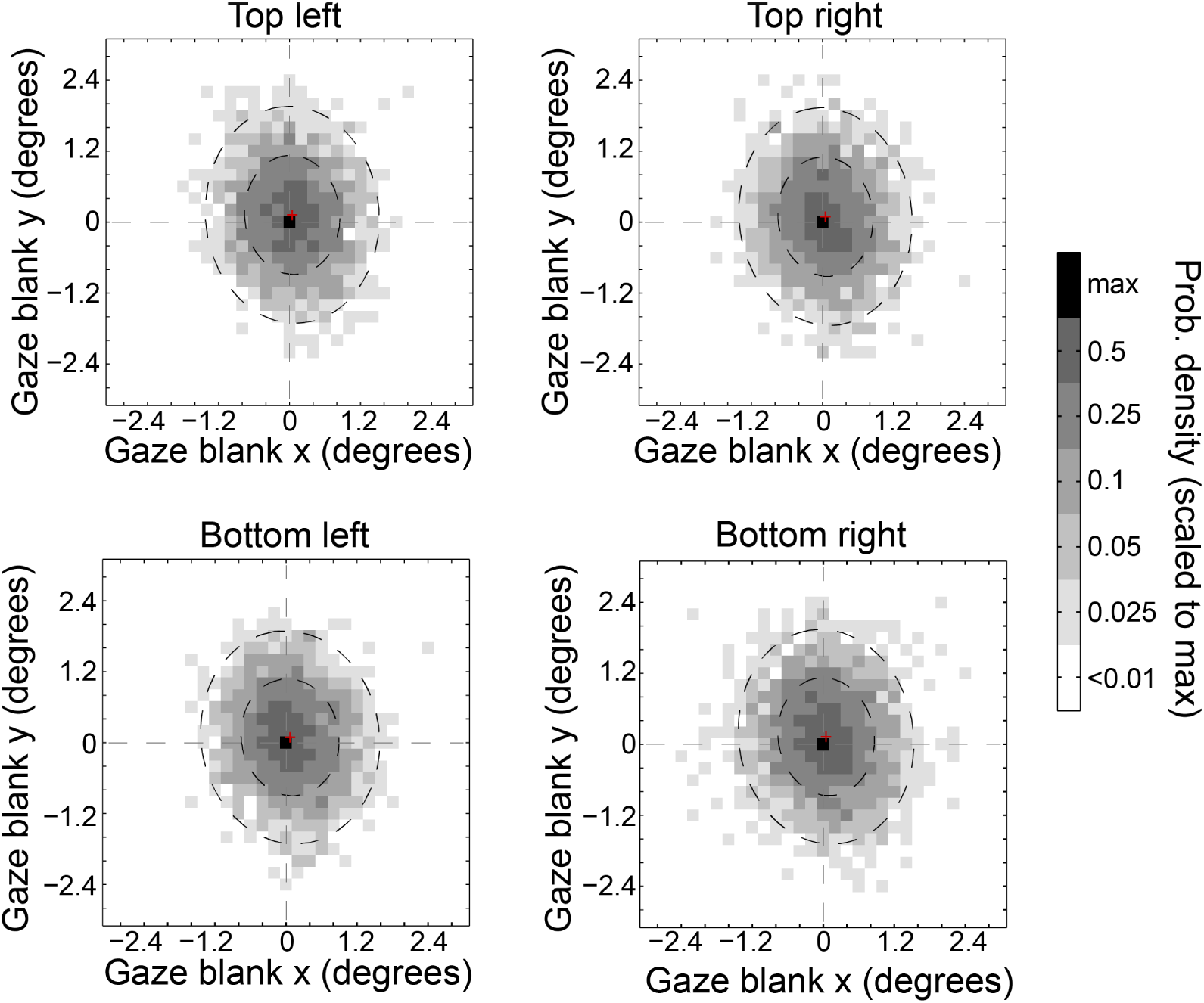
Fixation location during last 10ms of pre-target blank screen in 4-quadrant extension. To present the effect of stochastic context, fixation locations are presented as function of the most likely target location in HP condition. Densities were calculated in 0.1 × 0.1 degrees^2^, merging data points from all participants and normalizing to the maximum value per condition. The single dark point marks maximal density and is always at the screen center; red crosses indicate mean values for condition; inner/outer circles mark areas encompassing 50% and 90% of all fixations.

### Bootstrap ΔFB

As reported in the main text, we considered the quantity Δ*FB* = *mean* (*FB*pret70) – *mean* (*FB*pret30) as a measure of sensitivity to global statistics. However this grand-average quantity could also reflect the different proportion of alternate and return trials in the two conditions: given that return trials induced positive FB (in both conditions), a greater proportion of returns could bias the overall statistic even if returns had the same impact on FB in both conditions. This concern only applies to the grand-average measure; other analyses that quantified trial-by-trial effects or partialed out the impact of last transition do not share it. To evaluate this issue we used boostraping to create surrogate bootstrapped series, for each condition, so that each contained an equal number of alternate and return trials, and evaluated ΔFB in those.

These were constructed as follows. For each participant, we counted the number of alternate trials in the pret70 condition (*nalt*_70_). We then generated 100 surrogate distributions of 2 × *nalt*_70_ elements with all the elements sampled from the pret70 condition: *nalt*_70_ elements were sampled with replacement from the return trials and *nalt*_70_ elements were sampled with from the alternate trials. This produced 100 *bootFB*_*pert*70_ distributions. Similarly we calculated the number of returns in the pret30 condition (*nret*_30_) and we derived 100 *bootFB*_*pert*30_ distributions with an equal number of alternate and returns. We could then derive ΔFB for these bootstrapped series as in the main analysis, *boot*Δ*FB* = mean *bootFB_pret70_* – mean *bootFB_pret30_*. Averaging across participants we obtained mean *boot*Δ*FB* = 0.27 ± 0.03° in the first 50 trials and mean *boot*Δ*FB* = 0.39 ± 0.04° in the second half of trials. These values were significantly different with *t*(20) = 4.48, *p* < .001, *d* = 0.92. This analysis shows that it is the distribution of values of alternate and return trials that drives ΔFB rather than the proportions of the two types of trials.

**Figure AI.3.**
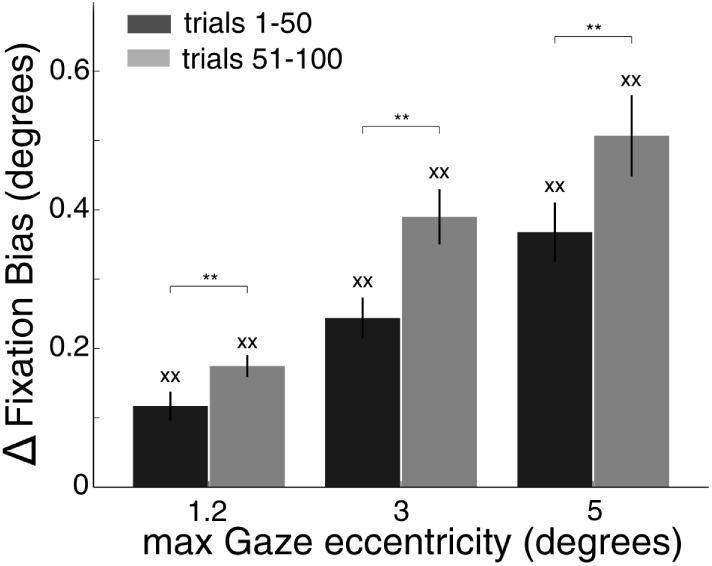
Δ*FB* calculated in first half of trials (dark gray bars) and in the second half of trials (gray bars) with three different limits to the admitted FB. Crosses above each bar indicate significant differences from zero. Asterisks above bar pairs indicate significant differences.

### RW-model validation

To evaluate the validity of the RW models, we used a leave-one-series-out validation scheme on the single-participant level. For each condition, we fit the model parameters from nine of the ten series, and the resulting parameter set was then evaluated against the left-out series. Specifically, model-derived series were generated by applying the updating scheme of Equation 2 to the true sequence of screen side transitions in the left-out series. Since every session was left out once, and the left-out time series could have a different number of valid trials in each fold, we evaluated the goodness of fit (percentage of explained variance) for each series using the adjusted coefficient of determination, 
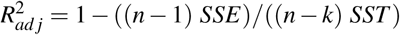
, where *n* is the number of points of the validating session and *k* is the number of free parameters; *SSE* is the sum of squared fit errors and *SST* is the sum of squared deviation from the mean of the series to predict. The reported variance reduction per participant was the mean of the adjusted coefficients of determination calculated for each of the ten validations. To determine whether the individuals’ variance reductions were significantly greater than would be expected by chance, we constructed synthetic series of alternations and returns that predict, through the estimated RW model parameters, the left-out FB data. This was done by permuting the sequence of screen side transitions in the left out series (1000 times). The participant’s mean variance reduction was than ranked in relation to the mean distribution of the permuted variance reduction.

### Mutual Information

We used MI to quantify the amount of information that is conveyed by FB and SL about the overall statistic of the target locations. Since SL on any given trial depend on whether it is an alternate or a return (due to IOR) we considered this factor in the MI calculation and computed the quantity *I(pret; SL\y)*, where y just defines if the trial is an alternate or return (Equation 7)

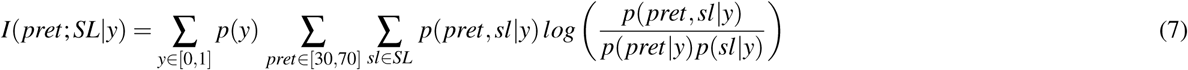

We calculated all MI quantities and their bias correction through the ‘Gaussian method’, (i.e. considering the probability distributions as Gaussians) (Misra, Singh, & Demchuk, 2005). Calculation using the ‘direct method’ gave similar results (Magri, Whittingstall, Singh, Logothetis, & Panzeri, 2009). In both cases we performed a bootstrap correction to reduce upward bias (Panzeri et al., 2007).

### LATER model applied to saccade latencies

The LATER model of saccade latency (Carpenter & Williams, 1995) treats SL as the time needed for a linear evidence accumulator to reach threshold. The accumulation rate (*r*) is considered to be normally distributed (mean rate μ and standard deviation σ) and these are the only independent parameters of the model. Nevertheless as implemented in prior studies, it is useful to explicitly derive a threshold parameter (*θ*), since it may reflect the a-priori state before the appearance of the target (Noorani & Carpenter, 2016). We applied this model to our SL data, first dividing trials according to last transition, where *r_j_* = *N* (μ,σ).:

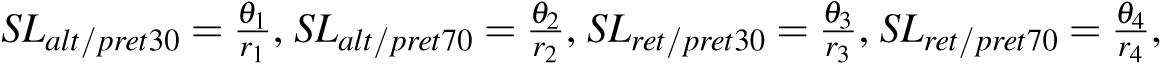

For each condition and for each participant, we then estimated the parameters *θ* and μ by minimizing the likelihood function, 
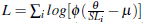
 where ϕ is the standard normal probability density function (following Kim et al., 2017). Each parameter was then normalized by its mean across different conditions. In five participants we excluded a small subset (about 5%) of trials whose values on the recinormal plot lay on a line with a smaller slope respect the other points suggestive of express saccade dynamics (see Carpenter, 1994).

## Extension to 4 quadrants

### Stimuli generation

To equally represent all the possible eye trajectories, in the highly predictive condition we generated series of target locations using four different transition matrices. When considered across these four matrices, the proportion of pair-wise transitions was equal for the HP and LP processes. This meant that statistical structure was not confounded with the pair-wise movements across trials. Specifically, in addition to the matrix *M_HP1_* presented in the main text, we used also *M_HP2_* below, and two transpositions of those matrices, 
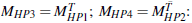

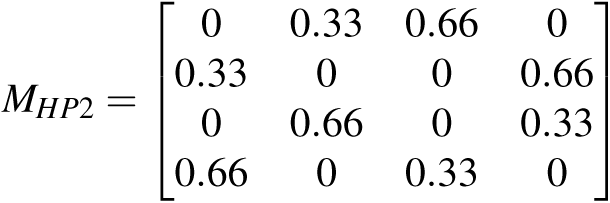

Given these four instantiations of the HP process, the LP matrix (in the main text) is the average of the HP matrices. Each matrix (HP or LP) was used to generate two sessions, one with high predictable image category, one with low predictable image category. In all sessions we checked there was no mutual information between image category and image location.

We used four categories of RGB images with 125 different elements each. ‘Neutral faces’ taken from the Center for Vital Longevity Face Database (Minear & Park, 2004) and from a face database for machine learning studies (Vieira et al., 2014). All the other images were taken from public online repositories: ’fruits’, ‘musical instruments’ and ‘tools’. Within a category, the elements were randomly chosen with constraint that an image would not be shown on two consecutive sessions and never displayed in the same location.

### Temporal characterization of target location series

A property of Markov processes is that processes with different levels of transition probabilities have different periods of repetitions, which translates into different autocorrelation functions (and analogously, power spectra). Given the transition matrix *P*, its element *p_ij_* is the probability of a target appearing in location *l_j_* given that on the current trial it is in location *l_i_*. Similarly the element *p_ij_* of the matrix *P^n^* = *PxP^n−1^* indicates the probability of having the target presented at location *l_j_* in exactly *n* transitions from its current location *l_i_*.

Consequently, for the HP matrices, the probability to complete the most likely transition after *n* transitions, by definition peaks at n=1 (p=0.66), but also after three trials (n=4, p=0.39) (Fig AI.4, panel A black circles). Conversely, for the LP matrix, the probability to complete an allowed transition after *n* transitions peaks at n=1 (p=0.33) and after two trials (n=3, p=0.26) (Fig AI.4, panel A gray diamonds). This recurrence characteristic of Markov processes underlie the temporal features of any possible allowed series of transitions. For instance we considered all the 2048 series of ten transitions a HP matrix allows and we calculated the expected autocorrelation in the x and y directions; we observed peaks at lag=3. Conversely, the autocorrelation of the series generated by the LP matrix, had a peak at lag=2. Finally we calculated the PSD of the series of target locations in the x-direction (see below), showing a peak for cycles of 3 trials in HP and of 2 trials in LP (the same held for the y-direction) (Fig AI.4, panel B).

### Spectral analysis of Gaze Bias

As input to the power spectra analysis we considered, separately, the horizontal (X) and vertical (Y) gaze coordinates of the single Gaze Bias measurement obtained in each trial, that is, the mean gaze position 10ms prior to target appearance. We did so because using the angle value provides less information about the strength of the anticipatory pattern (a single angle is consistent with multiple {*X,Y*} tuples), and because minor changes in X or Y would translate into large angular differences due to the relatively small deviations from center. The time series for this analysis were constructed by concatenating the 120 single trial Gaze Bias measures.

To overcome missing values, we first calculated the single-participant auto-correlation function of the gaze location over the prior 11 locations. R_xx_ (τ) = ∑*_k,(k – τ)∈[valid trials]_ x (k) x (k – τ)* where τ = 0,… 11 and *x* is the vector of Gaze Bias measurements in the x-axis. We constructed this vector by merging the values of all the sessions within a given condition (HP or LP), separated by a series of 12 *nan* values (as these were not valid trials). The power spectral density *S_x_* is then defined as the single sided Fourier transform of *R_xx_*. Similarly we obtained the cross-correlation function *R*xh* (τ)* and the cross spectrum *S_xh_ ( f)*, where *h* is the series of the horizontal coordinates of the target locations. Coherence in x-axis between the trial series and gaze position was defined as: 
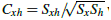
 where *h* is the screen side on which the target was located and *x* the Gaze Bias value (Mitra & Bokil, 2008). With the same procedure we obtained the spectrum *S_y_ ( f)* and cross spectrum *S_yv_ ( f)*, where *y* is the Gaze Bias in the *y* – *axis* and *v* is the vertical target location. Finally we obtained a measure of the total coherence as *C_Tot_* = *C_xh_* +*C_yv_*.

**Figure AI.4.**
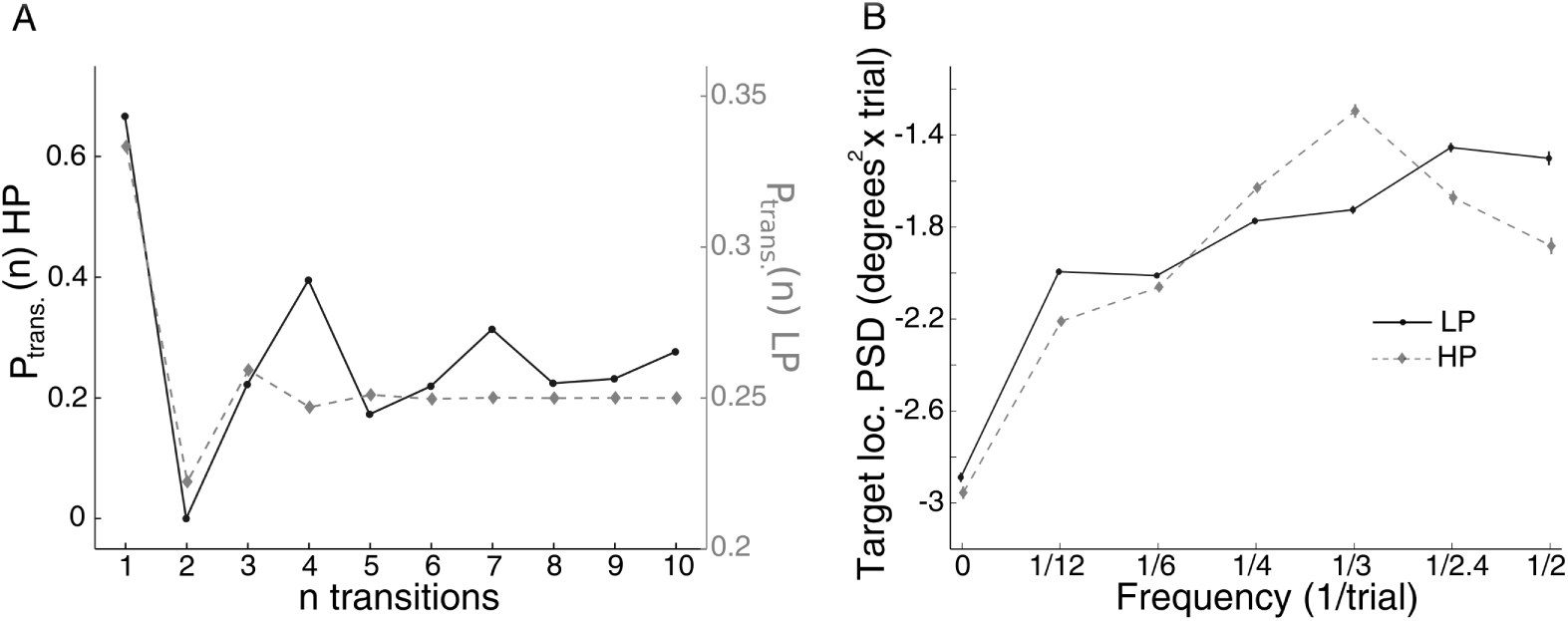
Recurrence properties of the experimental High Predictability and Low Predictability series. **Panel A**: Probability to complete the most likely transition after *n* steps in the highly predictive condition (HP; black circles) and to complete an allowed transition in the low predictability condition (LP; gray diamond). Data plotted on two separate axes. **Panel B**: Average PSD of target location series (x-direction) in HP (black circles) and LP (gray diamonds) conditions; only valid trials are included.

## Author contributions statement

G.N, D.M, and UH conceived the experiments, G.N. conducted the experiments, G.N and U.H analyzed the data. W.v.Z and D.M contributed methods. All authors reviewed the manuscript.

## Acknowledgements

We thank Leonardo Chelazzi for his comments. UH’s work was conducted in part while serving at and with support of the National Science Foundation. Any opinions, findings, and conclusions or recommendations expressed in this material are those of the author(s) and do not necessarily reflect the views of the NSF.

## Footnotes

1 We performed three validation and robustness analyses of Δ*FB*. First, we determined split-half reliability by deriving two separate Δ*FB* values per participant. one from odd trials and one from even trials. Split-half reliability was very robust (0.90 after correction). Second, we evaluated to what extent Δ*FB* depended on the specific trial inclusion criteria. We found that Δ*FB* was robust across a range of trial inclusion values, including trials where FB was restricted to 1.2° from screen center (see *Additional Information*). Third, we verified whether Δ*FB* was driven by transition structure or the number of returns and alternate trials in each series. We used bootstrapping to construct synthetic series from the pret70 and pret30 data, but where the number of alternation and return trials were equated (see *Additional Information*). We found statistically significant Δ*FB* values in these cases.

2 Group level t-tests of Beta values against zero. For pret70. (β_1_:*t*(20) = 7.40, *p* < .001, *d* = 1.61; β_2_:*t*(20) = 6.82, *p* < .001, *d* = 1.49; β_3_:*t*(20) = 3.39, *p* < .01, *d* = 0.74; β^4^:*t*(20) = 3.30, *p* = .01, *d* = 0.72; β_5_:*t*(20) = 2.72, *p* < .05, *d* = 0.59. for pret30. β_1_:*t*(20) = 7.34, *p* < .001, *d* = 1.60; β_2_:*t*(20) = 3.66, *p* < .01, *d* = 0.80; β_3_:*t*(20) = 4.70, *p* < .001, *d* = 1.03;. We note that for some lags, a few participants did show negative beta values for lags > 1; but there were only 18 such cases out of 147 beta values estimated.

3 The four Beta values were. β_1_:*t*(20) = 6.64, *p* < .001, *d* = 1.45; β_2_:*t*(20) = 5.33, *p* < .001, *d* = 1.16; β_3_:*t*(20) = 3.17, *p* = .01, *d* = 0.69; β_4_:*t*(20) = 4.41, *p* = .001, *d* = 0.96.

4 We conducted a parallel analysis for Gaze Bias Y (positive if in the upper screen side), considering as a third factor whether the previous target was on the upper or lower part of the screen; we did not find any significance in factors or interactions

